# Access to germinal center IL-4 microniches drives tissue-divergence of IgE memory responses

**DOI:** 10.64898/2025.12.03.692086

**Authors:** Sergio Villazala-Merino, Lucas Bertoia, Romain Fenouil, Myriam Moussa, Samuel Origlio, Claude Gregoire, Laura Almada, Mara Esposito, Stefano Colombo, Adriana Gruppi, Judith Allen, Andrew MacDonald, Nicolas Fazilleau, Carolyn G King, Mauro Gaya

## Abstract

Immunoglobulin E (IgE) drives allergic disease, yet how memory B cells (MBCs) reactivate to produce IgE, and how tissue localization shapes recall responses, remains unclear. Using mouse models of airborne exposure to house dust mites and *Alternaria*, we found that allergen sensitization generates lymphoid- and lung-resident MBCs. Upon allergen re-exposure, these populations followed distinct differentiation trajectories: lymph node MBCs engaged a germinal center (GC)-dependent pathway that generated both IgG1⁺ and IgE⁺ plasma cells (PCs), whereas lung MBCs followed a GC-independent route producing mainly IgG1⁺ PCs. GC re-entry granted MBCs access to an IL-4-rich microniche formed by Tfh cells, which was essential for IgE production. Disrupting GC re-entry, IL-4 signaling, or Tfh-derived IL-4 during recall markedly reduced allergen-specific IgE titers. These findings reveal a spatially and cytokine-restricted mechanism that confines IgE memory to lymphoid organs, positioning GC IL-4 microniches as anatomical safeguards against IgE production at barrier sites frequently exposed to environmental antigens.

## Introduction

During infection or immunization, B cells rapidly respond to antigens in secondary lymphoid organs, initiating processes that generate both immediate and long-lasting humoral immunity^1^. Activation induces expression of activation-induced deaminase (AID), which drives class switching to alternative antibody isotypes and fuels somatic hypermutation (SHM) within developing germinal centers (GCs)^2,3^. Within GCs, B cells iteratively refine their B cell receptor (BCR) affinity under the guidance of follicular helper T (Tfh) cells, while some cells with minimal affinity gain can also persist^4,5^. Successful GC B cells differentiate into two complementary long-lived populations: memory B cells (MBCs), which remain largely quiescent until antigen re-encounter, and long-lived plasma cells (PCs), which continuously secrete antibodies^6^. Upon re-exposure, MBCs can rapidly generate antibody-secreting cells or, depending on the nature of the challenge, re-enter GCs to further optimize their BCR^7–9^. MBCs and long-lived PCs form a dual layer of humoral protection that ensures both speed and durability in immune defense^10^.

Systemic challenges lead to the accumulation of MBCs and long-lived PCs in primary and secondary lymphoid organs, with MBCs localized mainly in lymph nodes (LNs) and spleen, and long-lived PCs residing in the bone marrow^11,12^. In contrast, during local or mucosal infection, these populations are not confined to lymphoid tissues but can also establish residency in barrier sites exposed to the pathogen^13^. For instance, MBCs can persist in the lungs following influenza or SARS-CoV-2 infection, and long-lived PCs have been identified in the intestinal mucosa after *Salmonella* infection^14–16^. These tissue-resident populations exhibit limited recirculation, positioning immunological memory directly at sites of pathogen entry, where they enable rapid containment of infection and prevent systemic spread^17–19^. While this localization is advantageous for host defense, the presence of MBCs and PCs at barrier sites could become counterproductive in the context of innocuous antigens. Resident memory cells in these tissues may rapidly mount localized responses against harmless environmental triggers, such as pollen, house dust mites (HDM), animal dander, fungal spores, or certain foods, thereby contributing to allergic inflammation.

The main driver of pathogenic responses to allergens is immunoglobulin E (IgE), which binds to high-affinity FcεRI receptors on basophils and mast cells, priming them for future allergen encounters. Upon re-exposure, cross-linking of FcεRI-bound IgE by allergens triggers degranulation of these cells, releasing a cascade of pro-inflammatory mediators which contribute to tissue inflammation, vascular permeability, and smooth muscle contraction across affected organs^20^. The critical role of IgE in allergic diseases is underscored by the clinical efficacy of omalizumab, a monoclonal antibody that binds free IgE and prevents its interaction with FcεRI, thereby reducing allergic exacerbations and inflammation^21^.

In humans, allergen-specific IgE can persist in the circulation for extended periods even in the absence of recent antigen exposure, implying the existence of long-lived IgE-secreting PCs that maintain serological memory^22^. Supporting this notion, studies in mice have shown that chronic allergen exposure induces the generation of rare, long-lived IgE⁺ PCs in the bone marrow that continue to produce IgE and sustain allergic response^23,24^. Although these cells are uncommon, their secretion rates are exceptionally high, estimated to be at least five times greater than IgG1-secreting cells, allowing even a small pool to maintain clinically relevant IgE levels and drive chronic allergic inflammation^25^. Interestingly, recent evidence suggests that the spleen, rather than the bone marrow, can serve as a major reservoir for mature long-lived IgE⁺ PCs^26,27^.

Seasonal allergen exposures in humans can further trigger *de novo* waves of IgE production, suggesting that allergen-specific MBCs are reactivated upon re-exposure, renewing IgE secretion and amplifying allergic responses^28^. In line with these observations, recent studies in mouse and humans have identified an IL4rɑ⁺CD23⁺IgG1⁺ MBC subset that arises during allergic type 2 responses and harbors high-affinity clones capable of differentiating into IgE-producing cells upon allergen encounter^29,30^. However, despite the central role of this process in sustaining allergic disease, the mechanisms governing MBC differentiation into IgE⁺ PCs remain poorly understood. Moreover, whether MBCs residing in barrier tissues versus secondary lymphoid organs differ in their capacity to mount IgE responses upon allergen re-encounter remains elusive.

Here, we investigated the mechanisms underlying the differentiation of allergen-specific MBCs into IgE⁺ PCs and how tissue location shapes recall responses using mouse models of airborne allergen exposure, including HDM and *Alternaria*. Allergen sensitization generates both lymphoid- and lung-resident MBCs, and scRNA-seq combined with BCR-seq trajectory analyses revealed that MBC fate is strongly imprinted by anatomical location. Upon allergen re-exposure, LN MBCs follow a GC-dependent pathway, producing both IgG1⁺ and IgE⁺ PCs, whereas lung MBCs engage a GC-independent route, generating mainly IgG1⁺ PCs. Using inducible transgenic models that selectively interfere with recall responses while sparing primary responses, we show that GC re-entry grants MBCs access to an IL-4 microniche produced by Tfh cells, which is essential for IgE production. Collectively, these findings uncover a spatially and cytokine-restricted mechanism that confines IgE memory to lymphoid organs, limiting IgE responses at barrier sites frequently exposed to innocuous antigens.

## Results

### Allergen exposure seeds MBCs in lymphoid and lung compartments

To investigate B cell memory responses to an airborne allergen, we established an immunisation model based on intranasal administration of HDM extract. HDM is a ubiquitous environmental allergen derived from the *Dermatophagoides* species, and it is a major trigger of allergic rhinitis and asthma in humans^31^. Mice received one dose of HDM during the first week followed by five consecutive doses over the next two weeks (Figure 1A). Control mice were treated with PBS. We tracked the formation of GC (CD19⁺GL7⁺), PC (CD98⁺CD39⁺), and MBC (CD19⁺CD38⁺IgD⁻) populations in the mediastinal LN (mLN), lungs, and spleen by flow cytometry. To exclude circulating cells, we injected CD45 intravenously 5 minutes before sacrifice (Figure S1A).

**Figure 1.**
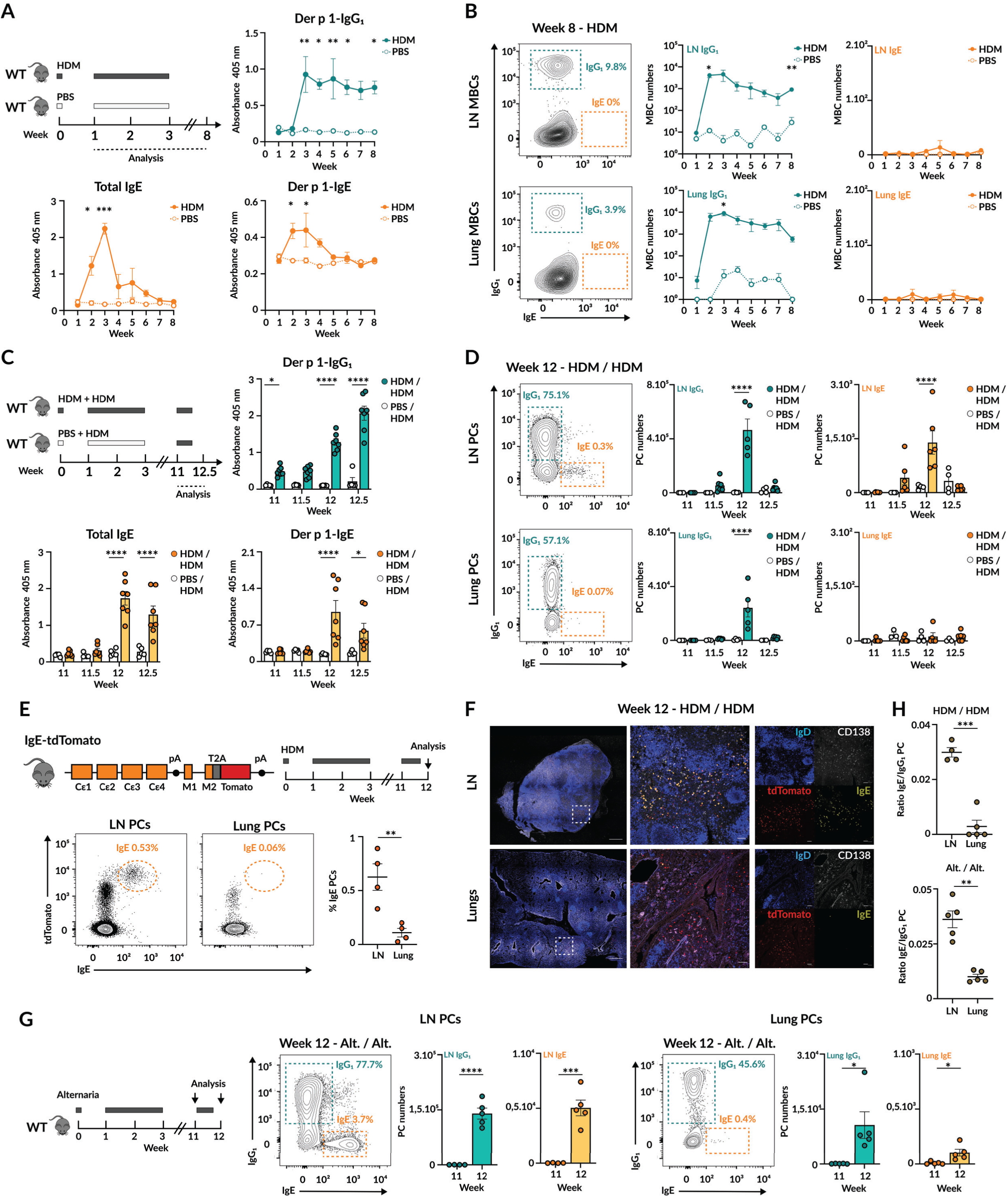
Tissue location dictates the ability of MBCs to mount IgE recall responses. (A) HDM immunisation model and experimental design (top left), Der p 1 IgG_1_ (top right), total IgE (bottom left) and Der p 1 IgE (bottom right) serum antibody titers. Dots correspond to mean ± s.e.m. n=3. (B) Contour plots show IgG_1_ and IgE MBCs in LNs and lungs after HDM immunisation along with IgG_1_ and IgE MBC cell number quantification. Dots represent mean ± s.e.m. n=3. (C) HDM immunisation and rechallenge experimental approach (top left), Der p 1 IgG_1_ (top right), total IgE (bottom left) and Der p 1 IgE (bottom right) serum antibody titers during HDM re-exposure stage. (D) Contour plots display IgG_1_ and IgE PCs in LN and lungs along with quantification of IgG_1_ and IgE PC numbers in both organs during HDM re-exposure. (E) Genetic construction of IgE-tdt Tomato mouse reporter (top left), experimental design (top right) and contour plot displaying Tomato^+^ and IgE^+^ PCs in LN and lungs (bottom) along with quantification of its frequency in each organ. (F) Confocal images of LN (top) and lung (bottom) sections from IgE-tdt Tomato mice treated with HDM as indicated in Figure 1E. Scale bars: left (500 µm), middle and right (50 µm). (G) *Alternaria alternata* (*Alt.*) immunisation and rechallenge model and experimental design (left) along with contour plot displaying IgG_1_ and IgE PCs in LN and lungs and absolute number of IgG_1_ and IgE PC in both organs during *Alternaria alternata* re-exposure. (H) Ratio of IgE to IgG_1_ PC in LN and lungs at week 12 after HDM re-challenge (top) and *Alt.* re-exposure (bottom) respectively. In all panels, quantification displays one representative experiment out of three. Unless otherwise indicated, each dot corresponds to one mouse and bars represent mean ± s.e.m. We used non-parametric multiple t-tests (A, B, C and D), nonparametric t-test (G) as well as paired t-test (F and H) for statistical analysis. * p<0.05, ** p<0.01, *** p<0.001 and **** p<0.0001.

We found that GC reactions developed in both mLN and lungs, peaking after three weeks of treatment, with GC activity gradually declining to baseline by week 7 in mLN but subsiding immediately after treatment termination in the lungs. In spleen, we found that GC reactions peaked two weeks after immunisation and then returned to background levels. Regarding PCs, we found that they emerged in mLN, lungs, and spleen, peaking two weeks after HDM exposure and returning to baseline by weeks 3-4 (Figure S1B). This indicates that most PCs were short-lived. To assess the specificity of the antibodies produced, we measured serum titers against Der p 1, the major HDM allergen. We found that Der p 1-specific IgG1 peaked at week 3 and remained stable throughout the observation period, likely reflecting the persistence of long-lived IgG1⁺ PCs in the bone marrow. In contrast, we observed that Der p 1-specific IgE and total IgE rose two weeks after immunisation, peaked one week later, and then progressively declined to baseline. This was in line with the notion that circulating IgE is captured by FceRI and Fcer2a, and that IgE⁺ long-lived PCs are not generated after acute HDM exposure but only after chronic treatment^23^ (Figure 1A). Importantly, we found that HDM immunisation induced the establishment of a long-lasting population of IgG1-expressing MBCs in secondary lymphoid organs and lungs, appearing after two weeks and remaining stable for at least eight weeks post-treatment (Figure 1B, S1C). Interestingly, we did not detect IgE⁺ MBCs in any organ tested.

Together, our results demonstrate that acute exposure to airborne allergens generates short-lived, allergen-specific antibody-secreting cells in secondary lymphoid organs and barrier tissues. However, the memory compartment in these organs is dominated by MBCs rather than long-lived PCs. This contrasts sharply with our recent findings in respiratory viral infections, where the memory landscape in lungs and draining lymphoid organs depends on the coordinated contribution of both MBCs and long-lived PCs^15^.

### Lymphoid organs are the primary sites of IgE⁺ recall responses

To assess whether MBCs residing in lungs and lymphoid organs can mount recall responses upon HDM re-exposure, we treated mice with either HDM or PBS as described in Figure 1A and allowed them to rest. At week 11, we challenged both groups with HDM extract (Figure 1C) and analyzed responses at weeks 11.5, 12, and 12.5. We found that in HDM-primed mice, re-exposure to HDM induced a rapid rise in Der p 1-specific IgG1 and IgE. IgG1 titers continued to increase over time, whereas IgE peaked one week after the first recall dose (Figure 1C). Importantly, we did not observe any upregulation of Der p 1-specific IgG1 or IgE antibodies in mice that were primed with PBS and only exposed to HDM during recall, consistent with the notion that rapid recall antibody responses depend on the presence of MBCs.

To pinpoint the tissue origin of secondary antibody responses, we measured PC responses in lymphoid organs and lungs. We found that short-lived IgG1⁺ and IgE⁺ PCs appeared in mLN and spleen one week after recall (Figure 1D, S1D). As hardly any IgE⁺ MBCs were detected after primary HDM exposure, these findings suggest that IgE⁺ PCs may arise from IgG1⁺ MBCs, as previously reported in humans^29,30^. In contrast, lung recall responses predominantly generated IgG1⁺ PCs, with very few IgE⁺ PCs, indicating that IgE⁺ PC differentiation is largely restricted to secondary lymphoid organs (Figure 1D). When we compared the relative magnitude of IgE⁺ and IgG1⁺ PCs, the IgE/IgG1 ratio confirmed a strong bias toward IgE⁺ PC development in lymphoid tissues (Figure 1H, top).

To validate these findings, we generated an IgEtdTomato reporter mouse by inserting a P2A sequence at the end of the last exon of the *Ighe* gene followed by a tdTomato reporter (Figure 1E). These mice were subjected to the same HDM priming, rest, and rechallenge protocol and analyzed at week 12. By flow cytometry, we identified IgE⁺ PCs as tdTomato⁺ cells co-expressing intracellular IgE. We found that tdTomato⁺IgE⁺ PCs were abundant in mLN but markedly reduced in lungs, confirming preferential IgE⁺ PC development in lymphoid organs (Figure 1E). We also observed an intermediate tdTomato⁺ population in both organs that did not express IgE. These cells may represent precursors poised to class switch to IgE under the right signals but appearing blocked in the lungs. To further characterize IgE-producing cells, we combined the reporter with an IgE-specific antibody that recognizes IgE expressed as a BCR but not IgE bound to FceR2a. Confocal microscopy revealed tdTomato⁺ cells co-stained with the IgE antibody predominantly in mLN and not in lungs, confirming active IgE expression in lymphoid organs (Figure 1F).

We next asked whether the preferential production of IgE recall responses in lymphoid tissues was strain dependent. To address this, we compared HDM recall responses in C57BL/6J and BALB/c mice. We found that both strains generated IgG1⁺ and IgE⁺ PCs in mLN, although BALB/c mice produced fewer IgE⁺ PCs than C57BL/6J. In lungs, both strains generated IgG1⁺ PCs but very few IgE⁺ PCs, which was reflected in a consistently reduced IgE/IgG1 PC ratio compared to mLN (Figure S1E). These findings suggest that the restriction of IgE⁺ PC development to lymphoid tissues is not strain dependent.

Finally, we tested whether this phenomenon extended beyond the HDM model by immunising and recalling mice with *Alternaria alternata* (Alt. alt.) extract using the same timeline as in Figure 1C. *A. alternata* is a ubiquitous airborne fungus and a clinically relevant allergen frequently associated with severe asthma^32^. We found that recall induced IgG1⁺ and IgE⁺ PCs in mLN and spleen, while lungs mainly sustained IgG1⁺ PCs with very few IgE⁺ PCs (Figure 1G, S1F). The IgE/IgG1 PC ratio again revealed a strong restriction of IgE⁺ PC development to mLN (Figure 1H, bottom).

Collectively, we found that recall responses consistently generate IgE⁺ PCs preferentially in secondary lymphoid organs rather than in lungs, highlighting the presence of a mechanism that enables IgE⁺ PC differentiation in lymphoid tissues in response to allergen re-exposure.

### MBCs give rise to IgE⁺ PCs in lymphoid organs upon recall

To determine whether the IgG1⁺ and IgE⁺ PCs arising after recall originate from MBCs, we performed fate mapping using the Aicda-CreERT2 Rosa26-EYFP model, which permanently labels AID-expressing B cells during HDM immunization at the time of tamoxifen administration (Figure 2A). Mice were treated with HDM and given tamoxifen between weeks 2 and 4, the peak of GC reactions. We assessed YFP labelling of MBCs at week 11 in mLN, lungs, and spleen. We found that YFP labelling was negligible in IgD⁺ naïve B cells, whereas 10–20% of MBCs were YFP⁺ across all three organs (Figure 2B, Figure S2A). Within the MBC pool, more than 40% of IgG1⁺ MBCs were labelled with YFP, compared to less than 20% of IgM⁺ MBCs, further validating the specificity of the model (Figure 2C, Figure S2B). In line with our previous observations (Figure 1B), no YFP⁺ IgE⁺ MBCs were detected in any organ.

**Figure 2.**
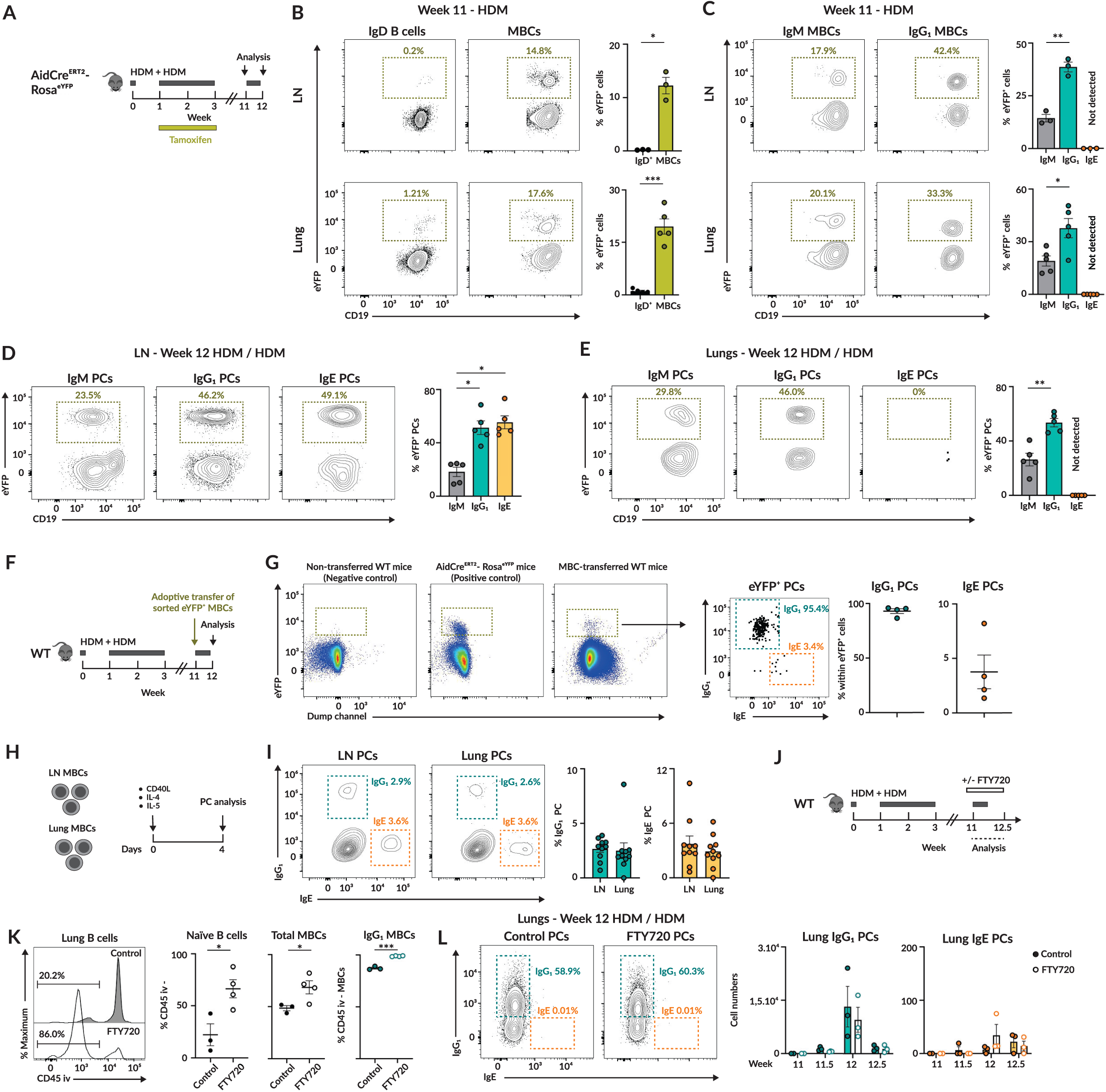
IgE recall responses originate primarily from lymphoid organ MBCs. (A) Experimental design for HDM treatment using AidCre^ERT2^-Rosa^eYFP^ mice. (B) Contour plots display YFP^+^ expressing B cells in IgD^+^ and MBC compartments in LN and lung. Bar plot shows quantification of the YFP^+^ frequency in each compartment. (C) Contour plots of YFP^+^ expressing B cells in IgM^+^, IgG ^+^, and IgE^+^ MBCs in LN and lung. Bars represent frequency of YFP^+^ cells for each isotype of MBCs. (D) Contour plots of YFP^+^ expressing B cells in IgM^+^, IgG ^+^, and IgE^+^ PCs in LN. Bars correspond to the percentage of YFP^+^ cells present within each PC population. (E) Contour plots of YFP^+^ expressing B cells in IgM^+^, IgG ^+^, and IgE^+^ PCs in lungs. Bar plot represents the quantification of the frequency of YFP^+^ cells for each isotype of PCs. (F) Experimental design for YFP^+^ MBC adoptive transfer (sorted following gating strategy depicted in Figure S1A). (G) Pseudocolour plots display YFP^+^ labelling (gated in CD19^+^ YFP^+^ cells) and dot plot showing IgG_1_ and IgE PCs (gated on YFP^+^ CD39^+^ CD98^+^) in lymphoid organs including quantification of their contribution to IgG_1_ and IgE PCs. (H) *In vitro* culture conditions of sorted LN and lung MBCs (sorted following gating strategy depicted in Figure S1A). (I) Contour plots show IgG_1_ and IgE PCs differentiated in vitro from LN and lung MBCs including quantification. (J) Experimental design for FTY720 treatment. (K) Histogram shows B cell labelling with anti-CD45 antibody administered i.v. along with quantification of naïve (IgD^+^) B cells (left), MBCs (middle) and IgG_1_ MBCs (right) protected from staining in lungs of untreated and FTY720 treated animals. (L) Contour plots display IgG_1_ and IgE PCs in lungs of untreated and FTY720 treated animals at week 12. Quantification of lung IgG_1_ and IgE PCs during HDM re-exposure. In all panels, quantification displays one representative experiment out of three. Each dot corresponds to one mouse and bars represent mean ± s.e.m. We used one-way ANOVA (B, C, D and E), paired t-tests (H) and nonparametric multiple t-tests (K and L) for statistical analysis. * p<0.05, ** p<0.01 and *** p<0.001.

We next examined the PC compartment upon HDM recall. In both mLN and spleen, we observed that approximately 25% of IgM⁺ PCs were YFP⁺, whereas around 50% of IgG1⁺ and IgE⁺ PCs were YFP⁺ (Figure 2D, S2C). These findings support the notion that, upon HDM recall, IgE⁺ PCs in lymphoid organs arise from previously labeled memory cells. In line with our previous results, in the lungs we observed YFP⁺ labeling in ∼30% of IgM⁺ PCs and ∼50% of IgG1⁺ PCs, but very few IgE⁺ PCs were detected, and none of them were YFP⁺ (Figure 2E). To unequivocally confirm that IgE⁺ PCs derive from MBCs and not from other YFP-labelled B cell populations, we performed an adoptive transfer experiment. YFP⁺ MBCs were sorted from mLN and spleens of HDM-immunised mice (gating strategy in Figure S1A), and 20,000 cells were transferred into competent hosts previously subjected to the same HDM immunisation regimen. Following re-exposure to five consecutive doses of HDM extract (Figure 2F), YFP⁺ cells gave rise to both IgG1⁺ and IgE⁺ PCs in all recipients (Figure 2G). Together, these results demonstrate that MBCs generated during the primary immune response perpetuate IgE responses during subsequent allergen exposure.

### LN and lung MBCs retain intrinsic potential to generate IgE⁺ PCs ex vivo

Given the scarcity of IgE⁺ PCs in the lungs after allergen re-exposure, we investigated whether lung MBCs are intrinsically limited in their ability to differentiate into IgE⁺ PCs. This limitation could result from the absence of a specific MBC subset capable of generating IgE⁺ PCs in the lungs, or from an intrinsic defect in lung MBCs that restricts their differentiation. To test this, we sorted MBCs from mLN and lungs of previously HDM-sensitized mice using the gating strategy in Figure S1A and cultured them ex vivo in a reductionist system. Cells were stimulated with CD40 ligand to mimic B-T cell interactions, and with IL-4 and IL-5, two cytokines abundantly produced during HDM exposure that favor IgE class switching and PC differentiation, respectively (Figure 2H). Four days later, we assessed MBC differentiation into IgG1⁺ and IgE⁺ PCs by flow cytometry. We found that mLN and lung MBCs differentiated to a similar extent into PCs (Figure S2D) and did so in comparable proportions into IgG1⁺ and IgE⁺ PCs (Figure 2I). These results indicate that lung MBCs are fully capable of differentiating into IgE⁺ PCs under permissive conditions, suggesting that the scarcity of IgE⁺ PCs in the lungs *in vivo* is not due to an intrinsic defect or absence of a specialized MBC subset.

Next, we considered whether the near absence of IgE⁺ PCs in the lungs could be due to their migration to lymphoid organs upon differentiation. To test this, we sensitized mice with HDM extract and treated them with FTY720, a drug that modulates sphingosine-1-phosphate receptor dynamics and blocks lymphocyte and plasma cell egress, starting one week before and during HDM recall (Figure 2J). Prior to recall, FTY720 treatment effectively blocked lung egress, increasing the proportion of B cells and IgG1⁺ MBCs resistant to intravenous anti-CD45 staining (Figure 2K). Upon HDM re-exposure, lung IgG1⁺ PCs developed in FTY720-treated mice similarly to controls. However, very few IgE⁺ PCs developed in lungs of FTY720-treated mice, comparable to untreated mice (Figure 2L). To ensure that FTY720 did not interfere with IgE⁺ PC development overall, we measured IgE⁺ PCs in mLN and observed no difference between treated and untreated mice (Figure S2E).

Together, these findings indicate that lung MBCs are not intrinsically incapable of differentiating into IgE⁺ PCs, nor are IgE⁺ PCs migrating away from the lungs. Rather, the absence of IgE⁺ PCs *in vivo* likely reflects an extrinsic difference in signals between lymphoid organs and lungs, with lymphoid tissues providing cues that promote IgE⁺ PC differentiation that are absent in the lung environment.

### Lung and LN MBCs adopt distinct differentiation paths upon recall

To explore why lung and LN MBCs differ in their ability to generate IgE⁺ PCs, we traced their differentiation trajectories during allergen recall using 5′-end single-cell RNA sequencing (scRNA-seq, 10x Genomics). Aicda-creERT2-Rosa26-EYFP mice were subjected to the HDM priming, rest, and rechallenge protocol, and YFP⁺ B cells were sorted from lungs and mLN at week 11 (pre-recall) and week 12 (one week post-recall) (Figure 3A). From each mouse, we collected up to 5,000 PCs and 8,000 MBC/GC B cells per tissue for downstream analysis.

**Figure 3.**
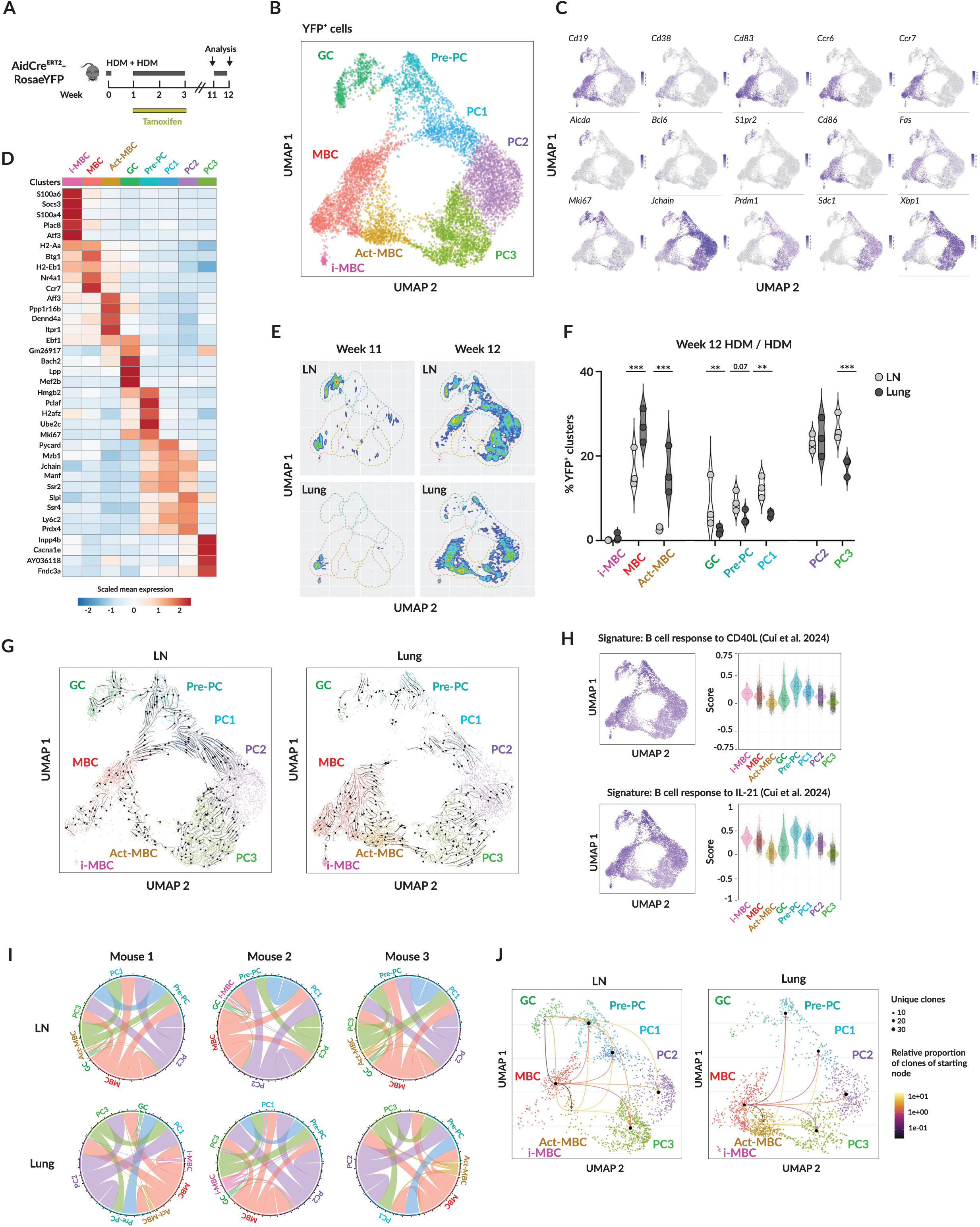
GC re-entry distinguishes LN from lung MBC recall responses. (A) Experimental design of single cell RNA sequencing (scRNA) using AidCre^ERT2^-Rosa^eYFP^ mice. (B) UMAP projection of YFP^+^ B cells in LN and lungs at week 11 and 12. (C) Feature plots show the expression of the indicated marker genes in YFP^+^ B cells laid out in the UMAP representation. Color denotes rate of expression. (D) Heatmap displays differentially expressed genes for each cluster. Colour indicates mean expression for each gene. (E) Density plots projected on the UMAP representation in LN and lungs separately at weeks 11 and 12. (F) Quantification of the contribution of YFP^+^ B cells to each cluster in LN and lungs at week 12. (G) RNA velocity analysis (scVelo) of YFP^+^ B cells in LN and lungs. (H) Module score for CD40L and IL-21 stimulated B cells (Cui et al, 2024) projected on UMAP (left) and quantified as violin plots (right) for each cluster. Color indicates expression rate. (I) Chord diagrams show clonotype sharing across clusters for LN and lung at week 12. (J) Representation of cluster interactions based on their clonal sharing probabilities for LN and lung. Three mice were used for each time and organ, mean ± s.e.m are represented. Each dot corresponds to one mouse. We used one-way ANOVA (F) for statistical analysis. ** p<0.01 and *** p<0.001.

Using dimensional reduction analysis with uniform manifold approximation and projection (UMAP), we identified eight distinct clusters of YFP⁺ cells (Figures 3B and 3C): three MBC clusters, one GC cluster, one pre-PC cluster, and three PC clusters. The MBC clusters were distinguished by expression of *Cd19*, *Cd38*, *Cd83*, *Ccr6*, and *Ccr7*. The GC cluster exhibited high expression of *Aicda*, *Bcl6*, *Cd86*, *S1pr2*, and *Mki67*. The pre-PC cluster showed elevated *Mki67* along with *Jchain*, *Prdm1* (Blimp1), *Sdc1* (CD138), and *Xbp1*, representing proliferating cells transitioning toward PCs. The remaining three clusters were designated as PCs due to high expression of *Jchain*, *Prdm1*, *Sdc1*, and *Xbp1*.

We then performed differential gene expression analysis to better characterize and identify each cluster (Figure 3D). Among the MBC clusters, the smallest cluster expressed *S100a6, Plac8* and lacked *Ccr6*, consistent with an ‘innate-like’ MBC (iMBC) population previously described^16^. The largest MBC cluster expressed antigen-presentation genes (*H2-Aa, H2-Eb1*) and regulatory genes such as *Btg1* and *Nr4a1* (Nur77), suggesting that it includes MBCs at steady state before rechallenge as well as those exiting GC reactions upon rechallenge. The third MBC cluster expressed transcriptional regulators (*Aff3*, *Ebf1*), genes linked to vesicle trafficking (*Dennd4a*) and mitochondrial maintenance (*Immp2l*), along with activation markers (*Cd83*, *Cd86*, *Fas*); because of this activated profile, we named this cluster Act-MBCs. The GC cluster expressed regulators of GC development and chromatin remodeling (*Bach2*, *Mef2b*, *Hmgb2*, *Pclaf*). The pre-PC cluster was enriched for DNA replication and cell cycle genes (*Hmgb2*, *Ube2c*, *Ptma*), reflecting its transitional state toward terminal PC differentiation. Among the PC clusters, PC1 and PC2 expressed genes linked to antibody production and PC differentiation (*Jchain*, *Manf*, *Mzb1*, *Prdx4*), whereas PC3 was enriched for genes involved in signaling and calcium flux (*Innp4b*, *Gphn*, *Cacna1e*).

To gain spatial and temporal information on the distribution of clusters, we separated YFP⁺ cells according to time (before and after rechallenge) and by organ (mLN and lung). At week 11, prior to re-exposure, YFP⁺ cells in mLN were found in MBC and residual GC clusters that were not actively producing PCs, whereas lung YFP⁺ cells were almost exclusively MBCs. Upon re-exposure, YFP⁺ cells differentiated into PCs in both organs. However, the Act-MBC cluster was highly enriched in lungs, while GC and pre-PC clusters were more frequent in mLN (Figures 3E and 3F). These results suggest that MBCs follow at least two distinct differentiation pathways upon allergen re-exposure: one that may involve GC re-entry leading to pre-PCs and PCs, and one that involves direct activation of MBCs into PCs. Notably, the contribution of these two MBC activation pathways upon recall is tissue-specific, with the pathway involving GCs dominating in mLN and the GC-independent, direct activation pathway prevailing in lungs.

To further resolve the differentiation dynamics among clusters, we performed RNA velocity analysis using scVelo (Figure 3G). In this analysis, arrows represent the predicted future transcriptional state of each cell, with their length reflecting the differentiation rate and their direction indicating the likely trajectory. In mLN, arrows indicated that MBCs predominantly transitioned through GC and pre-PC clusters before differentiating into PCs, with a minor GC-independent trajectory via Act-MBC. By contrast, lung MBCs almost exclusively followed the GC-independent pathway, transitioning through Act-MBC into PC2 and PC3. These results show that YFP⁺ B cells adopt distinct, organ-specific differentiation trajectories, with mLN favoring a GC-involving route and lungs favoring direct MBC activation.

To better characterize the pathways driving MBC differentiation, we overlaid a module score representing B cell gene signatures associated with responses to CD40L and IL-21, both abundant in the GC during B–T follicular helper cell interactions, onto the overall UMAP^33^. We found that cells with high CD40L and IL-21 module scores aligned predominantly along the GC-dependent differentiation pathway, highlighting the central role of GC-derived signals in shaping this trajectory (Figure 3H).

Because RNA velocity captures transcriptomic transitions but not lineage relationships, we next examined the clonal connectivity among clusters to better understand their origin and interactions. To this end, we performed single-cell BCR sequencing (scBCR-seq) on YFP⁺ B cells from mLN and lungs of mice re-exposed to HDM. Clonotypes were defined based on the edit distance of their CDR3 sequences using both light and heavy chain sequences, as previously described^34^. Pairwise clonal relationships between clusters were visualized using circos plots. Within each organ, we observed expanded clonotypes distributed across multiple clusters, confirming antigen-driven clonal responses. Importantly, MBCs displayed extensive clonal overlap with Act-MBC, GC, pre-PC, and all three PC clusters, strongly suggesting that MBCs seed the HDM recall response (Figures 3I and S3).

To quantify these relationships, we calculated clonal sharing frequencies to estimate transition probabilities between clusters. We found high clonal connectivity between GC, pre-PC, and all PC clusters, as well as between Act-MBC and PC clusters. These findings indicate that both GC-dependent and GC-independent (Act-MBC driven) pathways contribute to PC generation. Notably, clonotype sharing between GC and Act-MBC clusters was consistently absent in all mice analyzed, indicating that these populations represent mutually exclusive differentiation trajectories, consistent with the RNA velocity analysis (Figure 3J). This analysis also validated the tissue-specific preference of each pathway, showing that MBC responses in mLN are dominated by the GC-dependent axis, whereas in the lungs, the GC-independent route prevails.

Together, these findings demonstrate that MBC differentiation upon allergen recall is governed by tissue-imprinted programs, establishing anatomically segregated routes to PC formation.

### MBCs gain IgE fate through GC re-entry

Our scRNA-seq and scBCR-seq datasets revealed that MBCs can follow two distinct pathways to produce PCs: one involving GC reactions, which predominates in the mLN, and another proceeding through an intermediate Act-MBC population independently of GCs, which is more prominent in the lungs. Because our data indicated that IgE⁺ PCs arise from MBCs and are mainly produced in secondary lymphoid organs rather than in the lungs, we hypothesized that the GC-dependent pathway is required for IgE⁺ PC generation during recall responses. To test this hypothesis, we used Aicda-creERT2 Rosa26-EYFP mice and performed confocal imaging of mLNs before and after HDM re-challenge (Figure 4A). Prior to re-exposure, mLNs were small and contained few residual YFP⁺ GC structures with low Bcl6 expression and minimal PC production (Figure 4B). In contrast, after HDM recall, mLNs expanded nearly fivefold, displaying prominent GC areas containing YFP⁺ cells expressing high levels of Bcl6, as well as widespread YFP⁺ PCs expressing CD138 that occupied the medulla and T cell zones (Figure 4C). These results suggest that GC reactions could serve as a major site for the generation of IgE⁺ PCs upon HDM recall.

**Figure 4.**
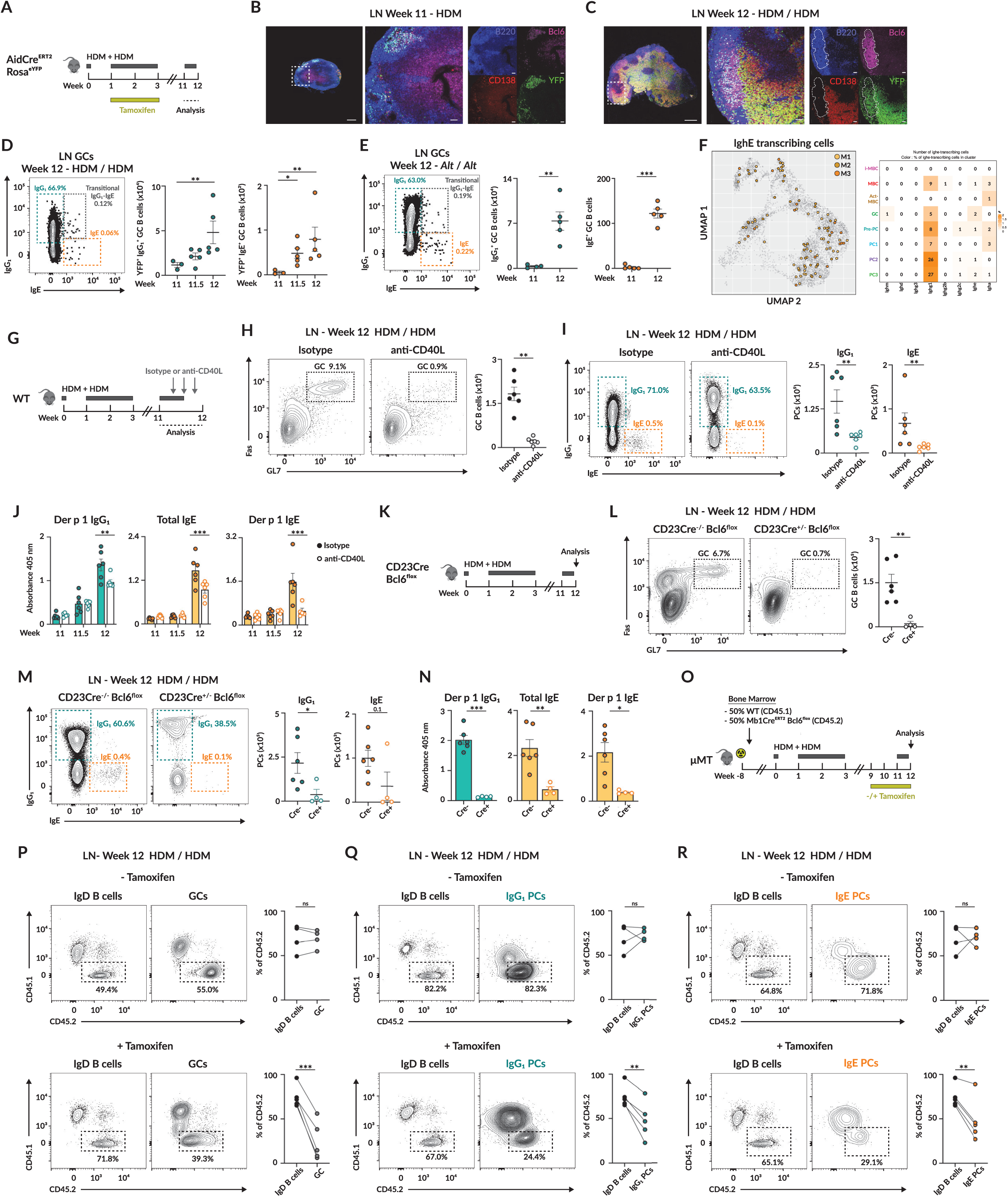
GC re-entry licenses MBCs for IgE production. (A) Experimental design for HDM treatment using AidCre^ERT2^-Rosa^eYFP^ mice. (B-C) Confocal images of LNs obtained at (B) week 11 and (C) week 12 from AidCre^ERT2^-Rosa^eYFP^ mice treated with HDM as in Figure 4A. Scale bars: left (500 µm), middle and right (50 µm). (D) Contour plot shows IgG1 and IgE YFP^+^ GC B cells in LN along with quantification following the HDM exposure protocol outlined in Figure 4A. (E) Contour plot shows IgG1 and IgE YFP^+^ GC B cells in LN along with quantification following *Alternaria* exposure protocol from Figure 1G. (F) LN YFP^+^ B cells expressing IghE transcripts projected in the UMAP shown in figure 3B. The table displays the absolute number of cells expressing other heavy chain genes and colour reflects percentage of IghE transcribing cells in each cluster. (G) Schematic of HDM sensitization, rechallenge and anti-CD40L administration. (H) Contour plots of GC B cells in LN of isotype and anti-CD40L treated mice. Absolute number quantification. (I) Contour plots of IgG_1_ and IgE PCs in LN of isotype and anti-CD40L treated animals along with cell number quantification. (J) Der p 1 IgG_1_ (left), total IgE and Der p 1 IgE serum antibody titers in isotype and anti-CD40L treated mice. (K) Diagram depicts experimental design for CD23^Cre^ -Bcl6^flox^. (L) Contour plots of GC B cells in LN of CD23Cre ^−/−^- and CD23Cre ^+/−^-Bcl6^flox^ mice along with absolute cell numbers. (M) Contour plots depict IgG_1_ and IgE PCs in LN of CD23Cre ^−/−^- and CD23Cre ^+/−^-Bcl6^flox^ mice including absolute cell number quantification. (N) Der p 1 IgG_1_ (left), total IgE (middle) and Der p 1 IgE (right) serum antibody titers in CD23Cre ^−/−^- and CD23Cre ^+/−^-Bcl6^flox^ mice. (O) Experimental approach describes mixed bone marrow chimera generation as well as HDM immunisation, re-exposure to HDM and protocol of tamoxifen administration for WT (CD45.1)/ Mb1Cre^ERT2^ Bcl6^flox^ (CD45.2) chimeric mice. (P) Contour plots exhibit CD45.1 and CD45.2 composition of IgD^+^ (left) and GC (right) in tamoxifen free (top) and tamoxifen fed (bottom) chimeric mice. (Q) Contour plots show CD45.1 and CD45.2 contribution to IgD^+^ (left) and IgG_1_ PCs (right) in tamoxifen free (top) and tamoxifen fed (bottom) chimeric mice. (R) Contour plots display CD45.1 and CD45.2 composition of IgD^+^ (left) and IgE PCs (right) in tamoxifen free (top) and tamoxifen fed (bottom) chimeric mice. In all panels, quantification displays one representative experiment out of three. Each dot corresponds to one mouse and bars represent mean ± s.e.m. We used one-way ANOVA (D, E and J), non-parametric t-tests (H, I, L and M) as well as paired t-tests (P-R) for statistical analysis. ns, non-significant; * p<0.05; ** p<0.01 and *** p<0.001.

To investigate this further, we measured IgG1 and IgE expression in YFP⁺ GC B cells before, during, and after HDM re-exposure in mLNs by flow cytometry. Before rechallenge, few IgG1⁺ GC B cells were detected in residual GCs, and their numbers increased rapidly after re-exposure, reaching a maximum one week post-rechallenge (Figure 4D). In contrast, no IgE⁺ GC B cells were detected prior to re-challenge; however, IgE⁺ YFP⁺ GC B cells progressively accumulated throughout the recall response. Notably, we identified a transitional YFP⁺ GC population co-expressing IgG1 and IgE after re-challenge, consistent with previous studies showing that IgE⁺ cells can arise via sequential class switching from IgG1⁺ precursors^35^. GC re-engagement was also observed following exposure to an independent airborne allergen, *Alternaria alternata*, resulting in accumulation of IgG1⁺ and IgE⁺ GC B cells and the emergence of a similar transitional IgG1⁺/IgE⁺ GC population (Figure 4E).

We next interrogated our scRNA-seq and scBCR-seq datasets to identify cells expressing *IghE* transcripts. The distribution of *IghE*-expressing cells aligned predominantly along the GC-dependent differentiation pathway in the UMAP, and most of these cells co-expressed *IghG1* transcripts, consistent with an IgG1-to-IgE sequential class-switching model (Figure 4F). Collectively, these results indicate that IgE⁺ PCs are likely generated within GC reactions upon allergen recall.

To experimentally assess this hypothesis, we disrupted GC responses during allergen rechallenge. To this end, we administered three doses of a blocking anti-CD40L antibody every two days during HDM re-exposure, thereby interfering with B-T cell interactions (Figure 4G). Importantly, treatment began two days post-allergen exposure to allow early B-T cell help while preventing later interactions in the GC. This intervention completely abrogated GC formation in the LN, lung, and spleen (Figures 4H and S4A). The absence of GC reactions led to a reduction in IgG1⁺ PC numbers in all three organs and a near-complete loss of IgE⁺ PC development in LN and spleen (Figures 4I and S4B). Correspondingly, total serum IgE titers were significantly lower, and no Der p 1-specific IgE was detected, whereas Der p 1-specific IgG1 was only partially reduced in anti-CD40L-treated animals (Figure 4J). These findings underscore a potential role of GC reactions in promoting MBC differentiation into IgE⁺ PCs.

To determine whether residual GCs present before re-challenge contribute to IgE⁺ PC production, we disrupted pre-existing GC reactions with a single dose of anti-CD40L administered two weeks prior to HDM re-exposure (Figure S4C). This treatment efficiently eliminated remaining GCs at week 11 (Figure S4D) and modestly reduced IgG1⁺ MBCs (Figure S4E). Upon HDM rechallenge, GCs re-formed in anti-CD40L-treated mice with similar kinetics, albeit to a lesser extent than in untreated animals (Figure S4F). Notably, IgE⁺ and IgG1⁺ GC B cells accumulated comparably in mice with disrupted pre-existing GCs and in untreated controls (Figure S4G). Collectively, these results indicate that MBCs can generate de novo GC reactions and give rise to IgE upon allergen recall.

To further validate the role of GC reactions in IgE responses, we subjected CD23Cre-Bcl6^flox^ mice, where Bcl6 deletion in B cells prevents GC formation, to the HDM priming, resting, and rechallenge protocol (Figure 4K). As expected, no GC structures were detected upon HDM re-exposure (Figures 4L and S4H). The absence of GCs resulted in a marked reduction of IgG1⁺ and IgE⁺ PCs across all organs analyzed following HDM rechallenge (Figures 4M and S4I). Consistent with these findings, neither total IgE nor allergen-specific IgG1 or IgE antibodies were detectable upon re-exposure (Figure 4N). These results support the notion that GC activity is required for the generation of memory responses to allergens.

However, because CD23Cre-Bcl6^flox^ mice are constitutively unable to form GC B cells, this model does not allow us to determine whether the impaired IgE⁺ PC response upon recall is due to defective MBC formation during primary immunization or to a failure of MBCs to engage the GC pathway upon recall. To overcome this limitation, we employed a tamoxifen-inducible model enabling Bcl6 deletion specifically in B cells during the recall phase. For this, irradiated B cell-deficient (μMT) hosts were reconstituted with a 1:1 mixture of wild-type (CD45.1) and Mb1Cre^ERT2^-Bcl6^flox^ (CD45.2) bone marrow cells to generate 50:50 mixed bone marrow chimeras. After eight weeks of reconstitution, mice were subjected to the HDM priming, resting, and rechallenge protocol. To induce Bcl6 deletion exclusively during the recall phase, tamoxifen was administered from week 9 onward, while control mice were kept tamoxifen-free (Figure 4O).

This competitive approach allowed us to assess the requirement of GC re-entry for IgE⁺ PC differentiation. In the absence of selective pressure, CD45.2⁺ cells (Mb1Cre^ERT2^-Bcl6^flox^) were expected to contribute equally to all B cell compartments. However, following Bcl6 deletion, we anticipated a selective reduction in CD45.2⁺ GC B cells and IgE⁺ PCs if GC participation was essential for IgE⁺ PC formation. As expected, chimeric mice not receiving tamoxifen displayed comparable proportions of CD45.2⁺ cells in both the IgD⁺ and GC compartments, whereas tamoxifen-treated chimeras showed a pronounced reduction in CD45.2⁺ GC B cells across all organs analyzed, confirming the efficiency and specificity of inducible GC disruption in our system (Figures 4P, S4J, and S4L).

Importantly, in the absence of tamoxifen, CD45.2⁺ cells displayed normal differentiation into IgG1⁺ and IgE⁺ PCs. In contrast, following Bcl6 deletion, the contribution of CD45.2⁺ cells to both IgG1⁺ and, more prominently, IgE⁺ PCs was strongly reduced relative to the IgD⁺ population in all organs (Figures 4Q, 4R, S4K, S4M and S4N). Together, these results demonstrate that GC reactions are required for effective IgE recall responses, supporting a model in which MBCs need to re-enter GCs to undergo terminal differentiation into IgE⁺ PCs.

### MBCs rely on GC IL-4 microniches to acquire an IgE fate

We next aimed to identify the signals received by MBCs along the GC-dependent differentiation pathway that instruct them to differentiate into IgE⁺ PCs. Allergen exposure induces high levels of type 2 cytokines, including IL-4 and IL-13. We interrogated our scRNA-seq dataset for expression of IL4rɑ, the common subunit of the IL-4 and IL-13 receptor. *IL4rɑ* was expressed at similar levels in MBC, Act-MBC, and GC clusters, indicating that B cells from both pathways are capable of sensing IL-4 and/or IL-13 (Figure 5A). Next, we asked whether B cells along the GC-dependent or GC-independent pathways show evidence of prior IL4rɑ engagement. Fcer2a, the low-affinity IgE receptor, is induced downstream of IL4rɑ signaling in B cells^16,36^. Analysis of our scRNA-seq dataset revealed preferential accumulation of *Fcer2a*-expressing cells in GC and MBC clusters, but not in the Act-MBC cluster, suggesting a potential role for IL-4/IL-13 specifically along the GC-dependent pathway (Figure 5B). To further assess cytokine exposure, we applied a module score based on genes induced by in vivo B cell exposure to individual cytokines^33^. Applying the IL-4-induced signature to our UMAP revealed higher scores along the GC-dependent differentiation trajectory compared with the GC-independent pathway (Figure 5C). No usable gene signatures were observed for IL-13 exposure, likely due to low IL13ra1 expression in naive B cells^33^. Together, these data suggest that sensing of type 2 cytokines along the GC-dependent pathway could play a role in directing MBC differentiation into IgE⁺ PCs.

**Figure 5.**
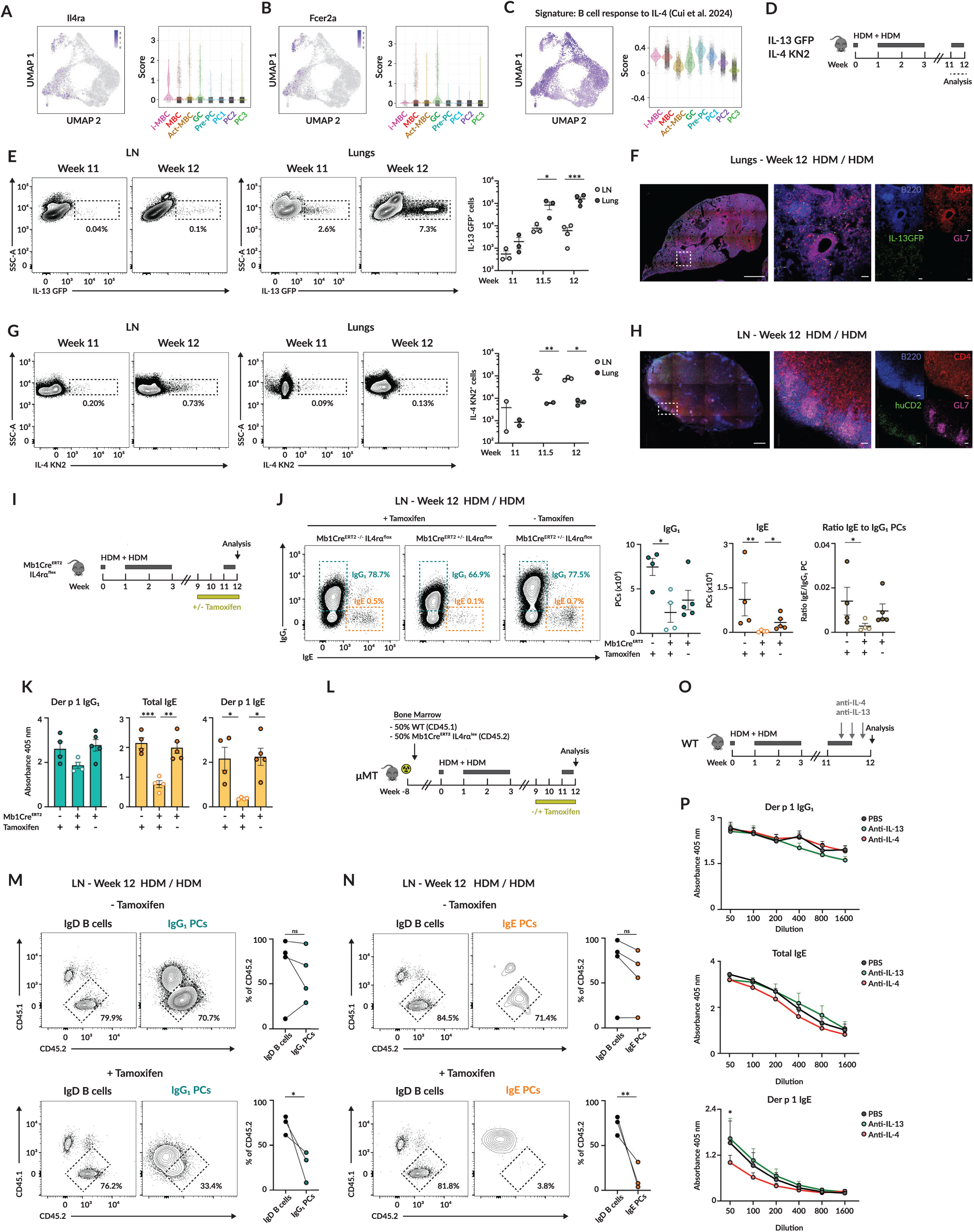
GC IL-4 microniches license MBCs for IgE differentiation. (A-B) Expression rate of (A) IL4rα and (B) FcεRI in YFP^+^ B cells projected on UMAP (left) and displayed on violin plots (right) for each cluster. (C) Module score of genes related to B cell stimulation with IL-4 (Cui et al, 2024) projected on UMAP and shown as violin plots for each cluster. (D) Schematic representation of experimental approach for IL-13 GFP and IL-4 KN2 mice. (E) Contour plots show IL-13 GFP expressing cells in LN and lungs including quantification of absolute cell numbers. (F) Confocal image of lungs from IL-13 GFP mice treated with HDM extract as indicated in Figure 5D. Scale bars: left (1000 µm), middle and right (50 µm). (G) Contour plots display IL-4 KN2 expressing cells in LN and lungs with absolute cell number quantification. (H) Confocal image of LN from IL-4 KN2 mice treated with HDM extract as indicated in Figure 5D. Scale bars: left (500 µm), middle and right (50 µm). (I) Experimental approach for HDM treatment and tamoxifen administration for Mb1Cre^ERT2^ IL4rα^flox^ mice. (J) Contour plots of IgG_1_ and IgE PCs in Mb1Cre^ERT2^ IL4rα^flox^ mice treated as in Figure 5I. Quantification shows the absolute cell numbers and IgE to IgG_1_ PC ratio. (K) Der p 1 IgG_1_ (left), total IgE (middle) and Der p 1 IgE (right) serum antibody titers in Mb1Cre^ERT2^ IL4rα^flox^ mice treated as in Figure 5I. (L) Representation depicting mixed bone marrow chimera generation, HDM treatment and tamoxifen administration regime for WT (CD45.1)/ Mb1Cre^ERT2^ IL4rα^flox^ (CD45.2) chimeric mice. (M-N) CD45.1 and CD45.2 contour plots on (M) IgD^+^ and IgG_1_ PCs as well as (N) IgD^+^ and IgE PCs in chimeric mice treated as in Figure 5L in the absence (top) or presence (bottom) of tamoxifen treatment. (O) Diagram describes experimental approach for IL-13 or IL-4 blockade. (P) Serum antibody titers of mice treated as in Figure 5O: Der p 1 IgG_1_ (top), total IgE (middle) and Der p 1 IgE (bottom). Dots represent mean ± s.e.m. n=3-4. In all panels, quantification displays one representative experiment out of three. Unless otherwise stated, each dot corresponds to one mouse and bars represent mean ± s.e.m. We used non parametric multiple tests (E, G, J and K), paired t-tests (M and N) and two-way ANOVA (O) for statistical analysis. ns, non-significant; * p<0.05; ** p<0.01 and *** p<0.001.

To gain insight into the spatiotemporal dynamics of IL-4 and IL-13 production upon recall, we subjected IL-13-GFP^37^ and IL4KN2^38^ reporter mice to the HDM priming, resting, and rechallenge protocol (Figure 5D). We observed that IL-13 production increased markedly in the lungs during HDM rechallenge, but was only modestly upregulated in the LN and spleen, approximately 10-fold lower than in the lungs (Figures 5E and S5A). Confocal imaging of IL-13-GFP lungs revealed that IL-13-producing cells were scattered throughout the tissue and largely co-stained with CD4, indicating that CD4⁺ T cells are the major contributors to IL-13 production during recall responses (Figure 5F). In contrast, IL-4 production was strongly upregulated in LN and spleen upon recall, with only modest increases in the lungs, approximately 10-fold lower than in the LN (Figures 5G and S5B). Confocal imaging further revealed that IL-4⁺ cells in LN clustered spatially with GL7⁺ GC B cells, suggesting that IL-4 production during recall is largely restricted to GC structures (Figure 5H). Together, these results demonstrate that type 2 cytokine production during HDM recall is anatomically segregated and highly compartmentalized: IL-13 is predominantly produced in the lungs by dispersed cells, whereas IL-4 is mainly produced in secondary lymphoid organs and is highly localized within GCs.

To examine the role of type-2 cytokines in IgE⁺ PC production during recall, we generated Mb1CreERT2-IL4rɑflox mice to inducibly delete the common IL-4/IL-13 receptor subunit specifically in B cells. These mice were subjected to the HDM priming, resting, and rechallenge protocol, and tamoxifen was administered only from week 9 onward to induce IL4rɑ deletion exclusively during the recall phase, leaving the primary response intact. Control mice were either Cre- mice treated with tamoxifen or Cre+ mice kept tamoxifen-free (Figure 5I). Upon HDM re-exposure, IL4rɑ deletion caused a modest reduction in IgG1⁺ PCs but almost completely abrogated IgE⁺ PC differentiation in LN and spleen (Figures 5J and S5C). In lungs, IgE⁺ PCs remained extremely rare regardless of IL4rɑ deletion, whereas IgG1⁺ PCs were modestly reduced (Figure S5D). In line with these results, Der p 1-specific IgG1 titers were slightly decreased, while total IgE and Der p 1-specific IgE titers were dramatically reduced when B cells were unable to sense IL-4/IL-13 during the recall phase (Figure 5K). To test whether this effect was B cell–intrinsic, we generated competitive chimeras by reconstituting μMT mice with a 1:1 mixture of WT (CD45.1) and Mb1CreERT2-IL4rɑflox (CD45.2) bone marrow. After eight weeks of reconstitution, mice underwent HDM priming, resting, and rechallenge. Tamoxifen was administered from week 9 onward to abrogate IL4rɑ signaling during the recall phase, while control mice remained tamoxifen-free (Figure 5L). In tamoxifen-free controls, CD45.2⁺ cells contributed equally to the IgD⁺ B cell, IgG1⁺ PC, and IgE⁺ PC compartments. In contrast, upon tamoxifen treatment, chimeras displayed a reduced contribution of CD45.2⁺ cells to the IgG1⁺ and, more prominently, to the IgE⁺ PC compartments relative to IgD⁺ B cells (Figures 5M and 5N). These results demonstrate that B cells require intrinsic IL4rɑ signaling for effective differentiation into IgE⁺ PCs.

Finally, to determine whether IL-4 or IL-13 is the key cytokine driving IgE⁺ PC development, we reduced IL-4 or IL-13 availability during HDM rechallenge by administering anti-IL-4 or anti-IL-13 blocking antibodies, respectively (Figure 5O). We found that IL-4 blockade during recall responses markedly reduced Der p 1-specific IgE serum levels, whereas IL-13 neutralization had no significant effect, with IgE levels comparable to those in control mice (Figure 5P). Collectively, these data identify IL-4 signaling as a critical checkpoint controlling the formation of allergen-specific IgE⁺ PCs upon recall.

### MBCs require Tfh-derived IL-4 to enter the IgE pathway

Given the marked organ-specific segregation of type 2 cytokine production upon allergen re-exposure, we investigated whether IL-4/IL-13 compartmentalization reflected differential distribution of T cell subsets across lymphoid organs and barrier tissues. To address this, WT mice were subjected to the HDM priming, resting, and rechallenge protocol, and the frequency of Th2 (GATA3^hi^) and Tfh (Bcl6⁺) cells was analyzed before and during rechallenge (Figure 6A). We found that HDM re-exposure led to an accumulation of GATA3^hi^ CD4⁺ T cells in both LN and lungs. In contrast, Bcl6⁺ Tfh cells were largely restricted to LNs, with very few detected in the lungs (Figures 6B and S6A). Imaging analyses confirmed these findings: in lungs, CD4⁺ T cells were scattered throughout the parenchyma with minimal infiltration into B cell clusters and no detectable Bcl6 expression, whereas in LN, CD4⁺ T cells infiltrated B cell follicles, co-expressed Bcl6, and localized within GCs (Figure 6C). To determine whether this compartmentalization extends to other allergens, we performed a similar analysis using *Alternaria* immunization and rechallenge (Figure 6D). As observed with HDM, Th2 cells developed across LN and lungs, with the highest abundance in the lungs, whereas Tfh cells were largely confined to LNs, comprising approximately 5–10% of CD4⁺ T cells (Figures 6E and S6B). Thus, T cell responses during allergen recall are spatially compartmentalized, with Th2 cells present in both lungs and lymph nodes but tending to accumulate more strongly in the lungs, particularly in response to *Alternaria*, whereas Tfh cells remain largely confined to secondary lymphoid organs.

**Figure 6.**
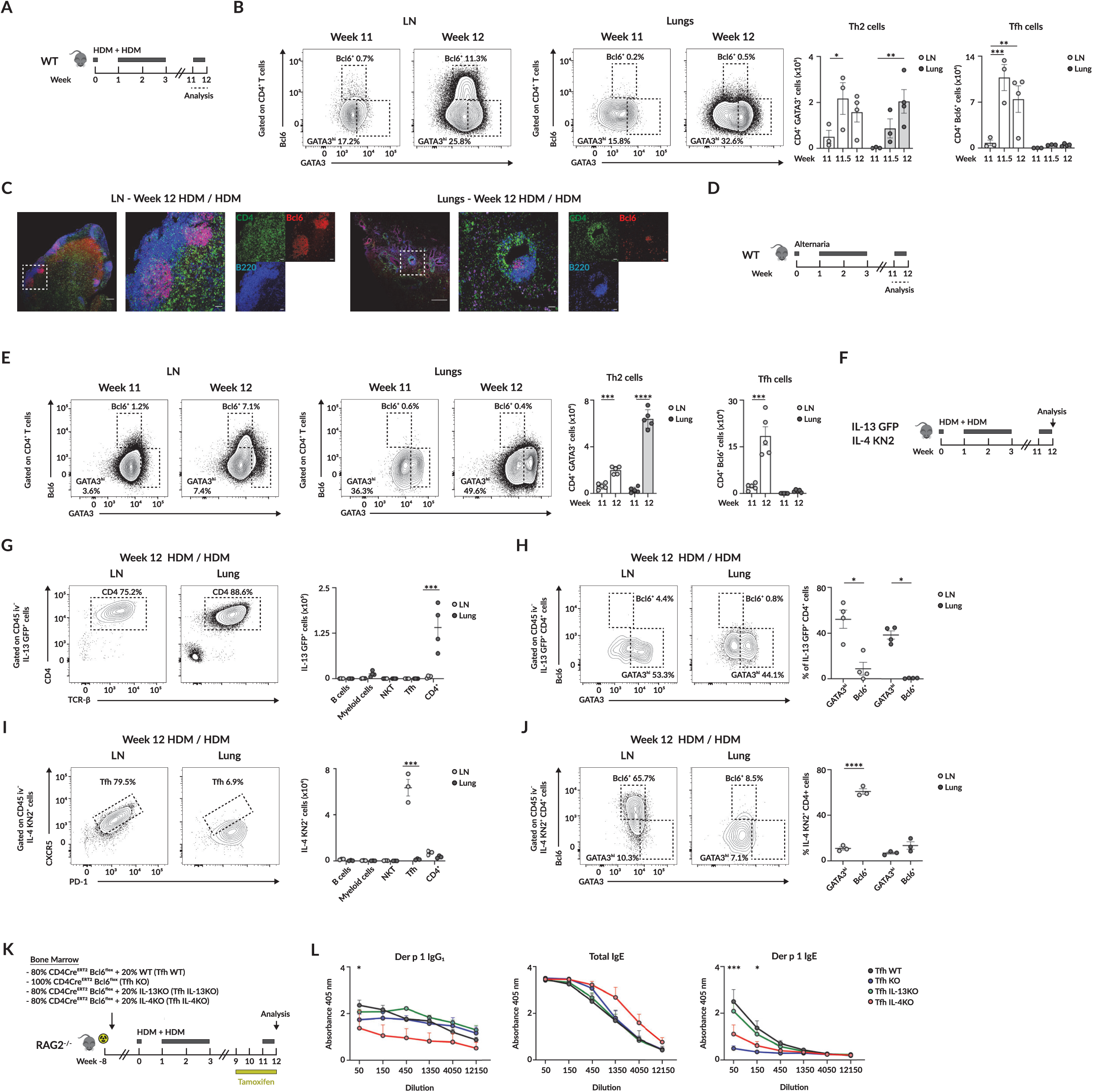
Tfh cells form GC IL-4 microniches essential for IgE recall. (A) Experimental approach for HDM treatment. (B) Contour plots display GATA3 and Bcl6 expression on CD4^+^ T cells in LN and lungs along with quantification of GATA3^hi^ and Bcl6-expressing CD4^+^ T cells. (C) Confocal images of LN (left) and lung (right) slides of mice treated with HDM according to protocol shown in figure 6A. Scale bars: left (200 µm for LN and 500 µm for lungs), middle and right (50 µm). (D) Experimental setup for *Alternaria* treatment. (E) Contour plots show GATA3 and Bcl6-expressing CD4^+^ T cells in LN and lungs including absolute cell numbers. (F) Schematics representing HDM treatment for IL-13 GFP and IL-4 KN2 mice. (G) Contour plots showing CD4 and TCR-β expression within IL-13-GFP expressing cells in LN and lungs. Quantification of IL-13-GFP^+^ expressing cells by cell type and organ. (H) Contour plots exhibit GATA3 and Bcl6 expression in IL-13-GFP^+^ CD4^+^ T cells in LN and lungs. Quantifications show the percentage of IL-13-GFP^+^ CD4^+^ T cells expressing GATA3^hi^ or Bcl6. (I) Contour plots display CXCR5 and PD-1 expression in the IL-4 KN2^+^ population in LN and lungs along with cell number enumeration by cell type and organ. (J) Contour plots exhibit GATA3 and Bcl6 expression in IL-4-KN2^+^ CD4^+^ T cells in LN and lungs. Quantifications show the percentage of IL-4-KN2^+^ CD4^+^ T cells expressing GATA3 or Bcl6. (K) Representation of bone marrow mixed chimeras construction as well as HDM treatment and tamoxifen administration to fully deplete Tfh (TfhKO) or to inducibly ablate IL-13 (Tfh IL-13KO) or IL-4 (Tfh IL-4KO) production exclusively by Tfh. (L) Serum antibody levels in mixed bone marrow chimeric mice shown in Figure 6K: Der p 1 IgG_1_ (left), total IgE (middle) and Der p 1 IgE (right). Dots denote mean ± s.e.m. n=5. In all panels, quantification displays one representative experiment out of three. Unless otherwise mentioned, each dot corresponds to one mouse and bars represent mean ± s.e.m. We used non parametric multiple tests (B, E), one-way ANOVA (G and I) paired t-tests (H and J) and two-way ANOVA (L) for statistical analysis. ns, non-significant; * p<0.05; ** p<0.01; *** p<0.001 and **** p<0.001.

We next examined whether this tissue-specific T cell distribution underlies the differential IL-4 and IL-13 production observed in lungs and LN. Using IL-13-GFP and IL-4-KN2 reporter mice subjected to HDM rechallenge (Figure 6F), we found that 60–90% of IL-13⁺ cells were CD4⁺ T cells across all organs, with markedly higher frequencies in the lungs compared with LN or spleen (Figures 6G and S6C). Approximately half of the IL-13⁺ CD4⁺ T cells co-expressed GATA3, confirming that Th2 cells represent a major source of IL-13 (Figures 6H and S6D). In contrast, IL-4-producing cells in LN and spleen displayed a canonical Tfh phenotype, characterized by CXCR5 and PD-1 expression, whereas the few IL-4⁺ cells detected in lungs lacked these markers (Figures 6I and S6E). Consistently, most IL-4⁺ cells in LN and spleen expressed Bcl6, confirming their Tfh identity, while the rare IL-4⁺ cells in lungs lacked both Bcl6 and GATA3 (Figures 6J and S6F). Together, these findings demonstrate that IL-13 secretion is primarily mediated by Th2 cells, predominantly in the lungs, whereas IL-4 production during allergen recall originates mainly from Tfh cells in secondary lymphoid organs.

To test whether IL-4 from Tfh cells is required for MBC differentiation into IgE⁺ PCs, we generated mixed bone marrow chimeras in which Tfh cells could be inducibly deleted or selectively deprived of IL-4 or IL-13 production during the recall phase. RAG2-deficient mice were reconstituted with 80% CD4Cre^ERT2^-Bcl6^flox^ bone marrow and 20% of either WT (Tfh WT), IL-4-deficient (Tfh IL-4 KO), IL-13-deficient (Tfh IL-13 KO), or CD4Cre^ERT2^-Bcl6^flox^ (Tfh KO) donor marrow. After eight weeks of reconstitution, mice were sensitized with HDM extract, administered tamoxifen from week 9 onward, and subsequently rechallenged (Figure 6K). Strikingly, Der p 1-specific IgE levels were markedly reduced in mice lacking Tfh cells or in which Tfh cells could not produce IL-4, whereas IL-13 deficiency in Tfh cells did not affect IgE titers compared with wild-type controls. (Figure 6L). These results demonstrate that IL-4 secreted by Tfh cells is essential for driving allergen-specific IgE during recall responses.

## Discussion

Our study reveals that the capacity to generate IgE during recall responses is not uniformly distributed across tissues but instead governed by the anatomical and cytokine landscape in which MBCs reside. Using physiologically relevant models of exposure to airborne allergens, we identify two distinct populations of allergen-specific MBCs: those maintained within lymphoid organs and those that persist in the lung tissue. Through transcriptional and clonal trajectory analyses, we find that this spatial segregation imprints fundamentally different recall behaviors. LN MBCs reacquire GC characteristics upon secondary challenge, accessing an IL-4-rich niche provided by Tfh cells that enables their differentiation into IgE⁺ PCs. In contrast, lung-resident MBCs follow a GC-independent differentiation route biased toward IgG1⁺ PC formation. Functional experiments using inducible genetic models confirm that GC re-engagement and Tfh-derived IL-4 are both indispensable for IgE recall. Together, these data establish that B cell memory to allergens is anatomically compartmentalized, with the ability to produce IgE confined to the structured environment of lymphoid GCs rather than barrier tissues. This spatial gating mechanism provides a previously unrecognized layer of control over humoral memory, limiting inappropriate IgE responses where antigen exposure is frequent.

Our initial observation that sensitization with HDM or *Alternaria* predominantly generates IgG⁺ MBCs, with very few IgE⁺ MBCs, is consistent with prior reports describing the scarcity of this population in both mice and humans^39,40^. This long-standing paradox has been attributed to the unique signaling properties of the IgE BCR, which differ markedly from other isotypes^41^. Several studies have demonstrated that engagement of the IgE BCR promotes rapid differentiation into antibody-secreting cells, thereby curtailing GC residency and precluding the formation of durable memory populations^42,43^. Moreover, IgE BCR signaling is characterized by chronic calcium flux and sustained activation of pro-apoptotic pathways, which further restrict the survival and maturation of IgE⁺ cells^44,45^. In addition, ligation of the IgE BCR on PCs can itself trigger apoptosis, highlighting the necessity of tight regulation over IgE expression to prevent aberrant activation^46^. Collectively, these mechanisms explain why most allergen-specific MBCs retain upstream isotypes such as IgG1 and rely on secondary class switching during recall to generate IgE⁺ PCs^35,47^.

Our data highlight a critical role for GC re-entry in granting MBCs access to IL-4 microniches and enabling their differentiation into IgE⁺ PCs. The capacity of MBCs to re-enter GCs during recall has been debated in the context of infection or immunization: some studies show that recall responses largely occur outside GCs, with limited MBC re-engagement, whereas others show that GC re-entry is possible and essential for generating high-affinity and broader responses^7–9^. In the context of allergic re-exposure, we used scRNA and BCR-seq and inducible deletion of GC B cells during recall to demonstrate that GC re-entry is indispensable for IgE⁺ PC differentiation. MBCs that bypass GCs, such as lung-resident populations, fail to access the IL-4-producing Tfh microniche and are unable to undergo sequential class switching to IgE. These results reveal that, in recall response to allergens, the GC functions not only as a site of affinity maturation but also as a spatial checkpoint that determines whether MBCs can acquire an IgE fate.

Our findings further emphasize the functional importance of IL-4 microniches within GCs. We previously showed that IL-4-producing NKT cells positioned at follicular borders initiate GC seeding during antiviral B cell responses^48^. In contrast, during recall to allergens, IL-4 production is confined to the GC core and derives exclusively from Tfh cells rather than NKT cells. Prior work, including ours, demonstrated that IL-4 availability within the GC regulates the output of antigen-specific memory B cells^16,49,50^. Here, we extend this paradigm by showing that confinement of IL-4 to GC microniches restricts IgE class switching to lymphoid tissues. This stands in sharp contrast to IgG class switching, which occurs earlier and often before GC entry^2^. Notably, the IL-4 microniche we describe parallels recently identified IL-2 microniches in LNs that drive Th2 cell differentiation from Tfh cells during allergen exposure, underscoring how spatial segregation of cytokines within lymphoid tissues underpins immune compartmentalization^51–54^.

In addition, our data reveal pronounced tissue polarization in cytokine production: IL-4 expression was confined to Tfh cells within lymph nodes, whereas IL-13 predominated in the lungs and was produced mainly by Th2 cells. This IL-4/IL-13 compartmentalization appears to be a general feature of mucosal type 2 responses, as similar patterns have been observed during helminth infection^55^. Recent studies have identified specialized Tfh13 cells that co-produce IL-4 and IL-13 to promote high-affinity IgE generation^56,57^. In our model, allergen-specific IgE titers were not significantly reduced in the absence of IL-13; however, we cannot exclude that IL-13 may influence the quality rather than the quantity of IgE, for instance by modulating affinity maturation or supporting the persistence of high-affinity clones. While our findings demonstrate that LN-resident MBCs can generate IgE upon recall by accessing IL-4-rich GC microniches, the contrasting cytokine landscape of the lung may impose additional constraints. Beyond lacking access to IL-4, lung MBCs are likely exposed to inhibitory signals such as TGF-β, which can antagonize IgE class switching^58^. Thus, tissue-specific cytokine organization not only provides the permissive cues for IgE differentiation in lymphoid organs but may also establish repressive environments in peripheral tissues. Future studies will be essential to determine how these opposing cytokine networks collectively shape the anatomical boundaries of IgE memory.

A recent study reported that lung-resident MBCs can locally produce IgE following repeated allergen exposure, which at first glance appears to contrast with our findings^59^. Their model used a single high-dose grass pollen sensitization, roughly tenfold higher than established protocols, followed by repeated ovalbumin challenges, creating unusually strong Th2 inflammation. In LatY136F mice, which carry a point mutation in the LAT adaptor and develop spontaneous and intense pulmonary Th2 responses with early lethality, we likewise observed local IgE production in the lungs^60^. Even in this setting, however, lymph nodes remained the dominant site of IgE generation, consistent with the idea that severe inflammation can relax but not fully erase normal anatomical constraints. In contrast, our approach incorporates sensitization, rest, and recall phases using physiologically relevant doses of airborne allergens such as HDM and Alternaria, providing a closer approximation to typical environmental exposures. Together, these observations suggest that IgE recall responses operate along an inflammatory continuum: under low-grade or physiological conditions, IgE production remains restricted to lymphoid organs, whereas in highly inflamed environments, strong local Th2 cues can partially override these constraints and permit local IgE generation.

Together, our findings outline a model in which IgE recall responses are anatomically and cytokine constrained. IgG1⁺ MBCs re-engage GCs to access IL-4-rich microniches, where they differentiate into IgE⁺ PCs This requirement distinguishes IgE recall from conventional memory responses, which often occur outside GCs. Such regulation, coupled with the spatial restriction of IL-4 to lymphoid GCs, acts as a safeguard that confines IgE production to controlled environments and prevents excessive or misdirected IgE responses at barrier surfaces. This compartmentalized organization may represent a broader feature of allergen-specific immunity, potentially extending to other barrier tissues such as the gut during food allergen exposure. Together, our study highlights how spatially organized cytokine cues and GC dynamics direct MBC fate during allergen exposure, ensuring IgE responses remain protective yet tightly controlled.

## Acknowledgements

We thank all the B cell Immunity to Infection lab for scientific discussions. We thank Pierre Milpied for advice on scRNA-seq. We thank Richard Locksley, Andrew McKenzie and Meinrad Busslinger for allowing us to use IL-4 KN2, IL-13 GFP and CD23-Cre mice, respectively. We thank Bernard Malissen for providing LatY136F mice, important to set up IgE staining protocols. We thank the NIH Tetramer Core Facility for the provision of labeled CD1d tetramers. We thank Claude-Agnès Reynaud and Jean Claude Weill for providing Aicda-CreERT2 Rosa26-EYFP mice. We thank Michelle Linterman for providing CD23-Cre mice. We thank Toby Lawrence and Nathalie Auphan for providing IL4ra flox mice. We thank the flow cytometry, imaging (Imagimm) and histology platforms for technical support and the biological resource unit for the breeding of animals (CIML).

## Author contributions

SVM and MG conceived the study, designed the experiments, analyzed the data, made the figures and wrote the paper. MG supervised the study. SVM, LB, MM, SO and CG performed the experiments. RF performed bioinformatic analysis. LA, ME, SC, AG, JA, AM, NF and CK provided materials. SC, NF and CK provided discussion.

## Data and materials availability

All data are available in the main text or the supplementary materials.

---

**Figure S1.**
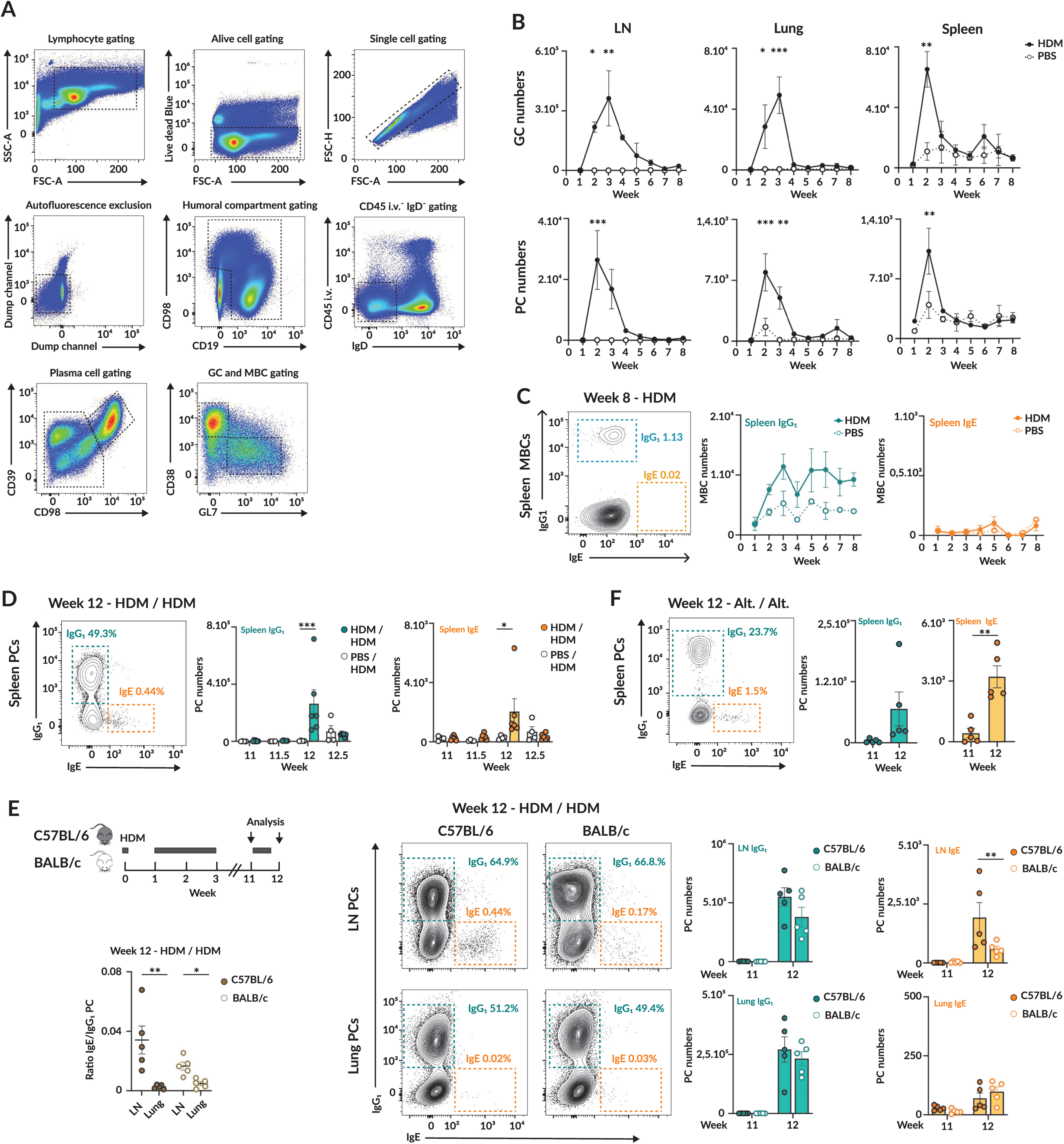
Tissue location dictates the ability of MBCs to mount IgE recall responses. (A) Gating strategy used for the identification of the different B cell populations. (B) Absolute cell number of GC and PC in LN (left), lungs (middle) and spleen (right) of mice immunized with HDM extract according to schematic in Figure 1A. Dots correspond to mean ± s.e.m. (C) Contour plots show IgG1 and IgE MBCs in spleen following HDM immunization as indicated on Figure 1A along with quantification of absolute cell numbers. Dots denote mean ± s.e.m. n=3. (D) Contour plots show IgG_1_ and IgE PCs in the spleen of mice treated with HDM extract as indicated on Figure 1C including absolute cell number quantification. (E) Diagram representation depicting HDM immunization and re-exposure to HDM in C57BL/6 and BALB/c mice (top left). Contour plots display IgG_1_ and IgE PCs in LN and lungs including absolute cell number quantification (right). Calculation of the ratio of IgG1 to IgE PC in LN and lungs in C57BL/6 and BALB/c mice (bottom left). (F) Contour plots display IgG_1_ and IgE PCs in the spleen of mice exposed to *Alt.* as shown in Figure 1G. Quantification shows absolute cell numbers. In all panels, quantification displays one representative experiment out of three. Unless otherwise indicated, each dot corresponds to one mouse and bars represent mean ± s.e.m. We used one-way ANOVA (B), non-parametric multiple t-tests (C, D, E and F) and paired t-tests (F, ratio IgE to IgG_1_ PCs) for statistical analysis. * p<0.05, ** p<0.01 and *** p<0.001.

**Figure S2.**
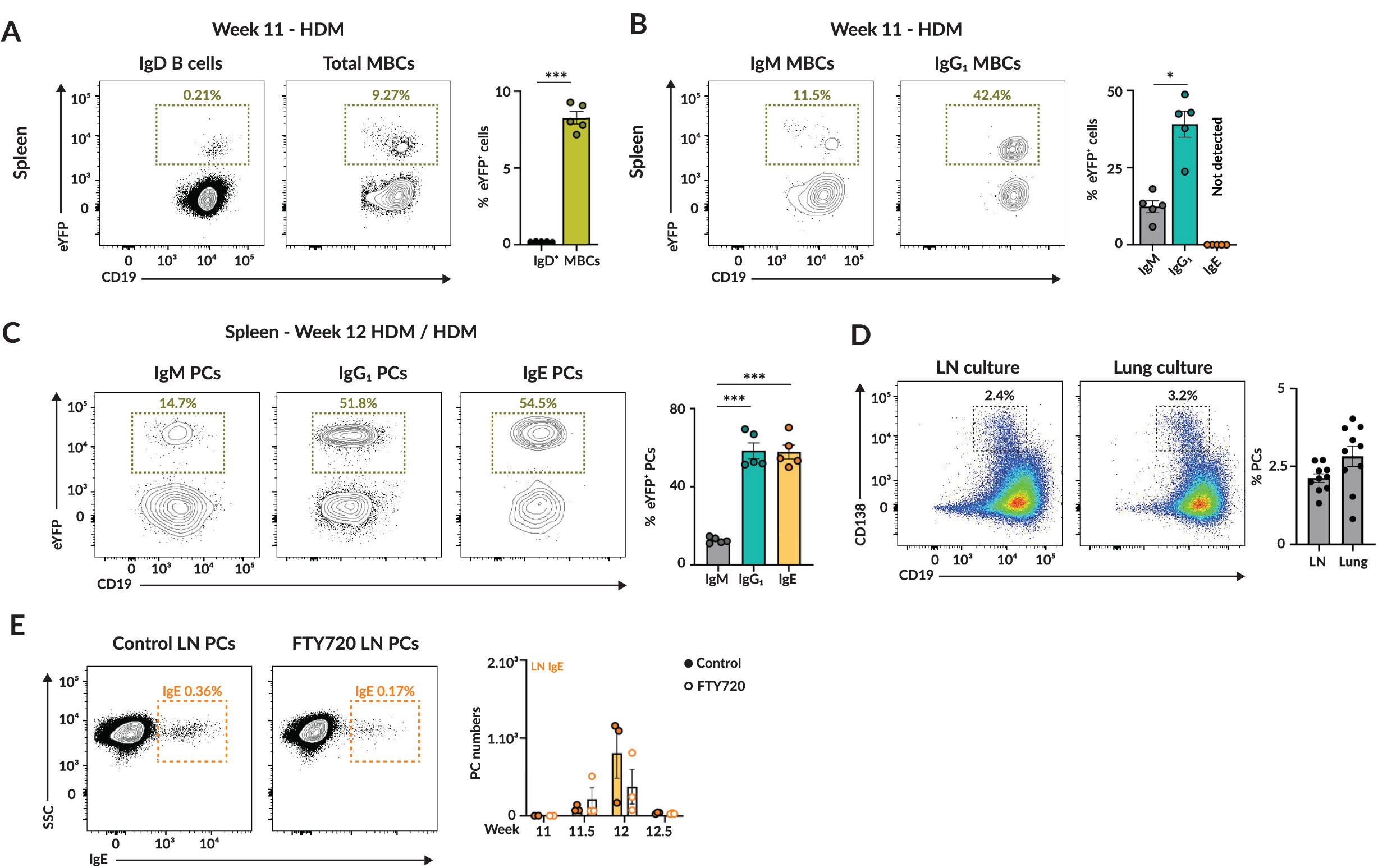
IgE recall responses originate primarily from lymphoid organ MBCs. (A-B) Contour plots compare YFP^+^ expression in (A) IgD^+^ and MBCs as well as (B) IgM^+^ and IgG_1_ ^+^ MBCs in the spleen of mice immunized with HDM extract as shown in Figure 2A including quantifications. (C) Contour plots show YFP expression within IgM, IgG1 and IgE PCs in the spleen of mice immunized and re-exposed to HDM extract as indicated on Figure 2A. (D) Pseudocolour plots display PC differentiation (CD138^+^) in *in vitro* stimulated LN and lung MBCs. Frequency of PCs in both cultures is shown. (E) Contour plots exhibiting IgE PC staining in LN of mice untreated or exposed to FTY720. Enumeration of IgE PCs is displayed. In all panels, quantification displays one representative experiment out of three. Each dot corresponds to one mouse and bars represent mean ± s.e.m. We used one-way ANOVA (A, B and C), paired t-tests (D) and nonparametric multiple t-tests (E) for statistical analysis. * p<0.05 and *** p<0.001.

**Figure S3.**
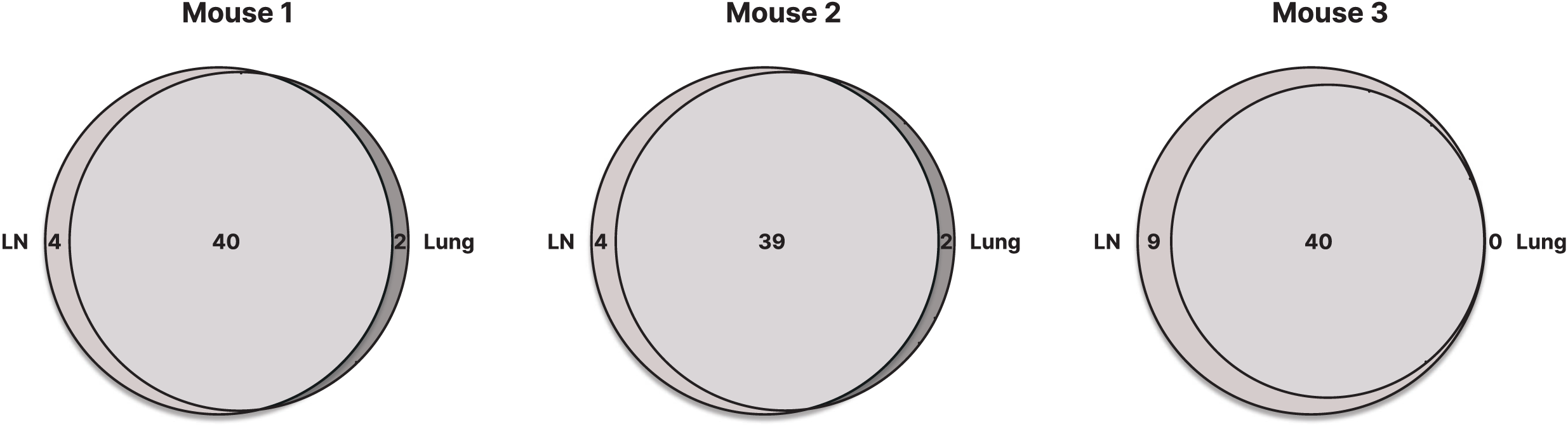
GC re-entry distinguishes LN from lung MBC recall responses. Venn diagrams present BCR clonotype overlap between LN and lungs for each of the three mice immunized and re-exposed to HDM extract as indicated in Figure 3A.

**Figure S4.**
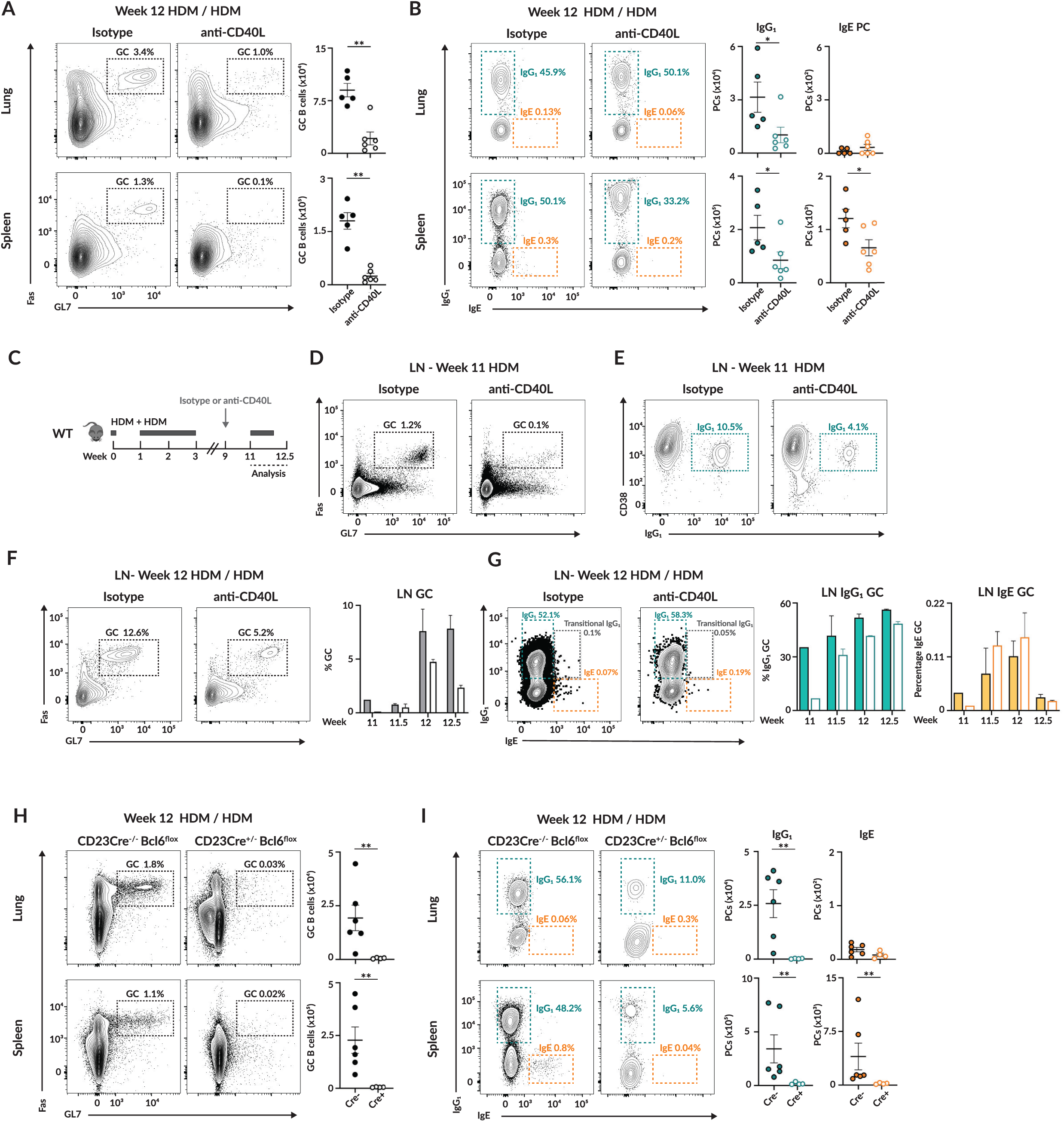
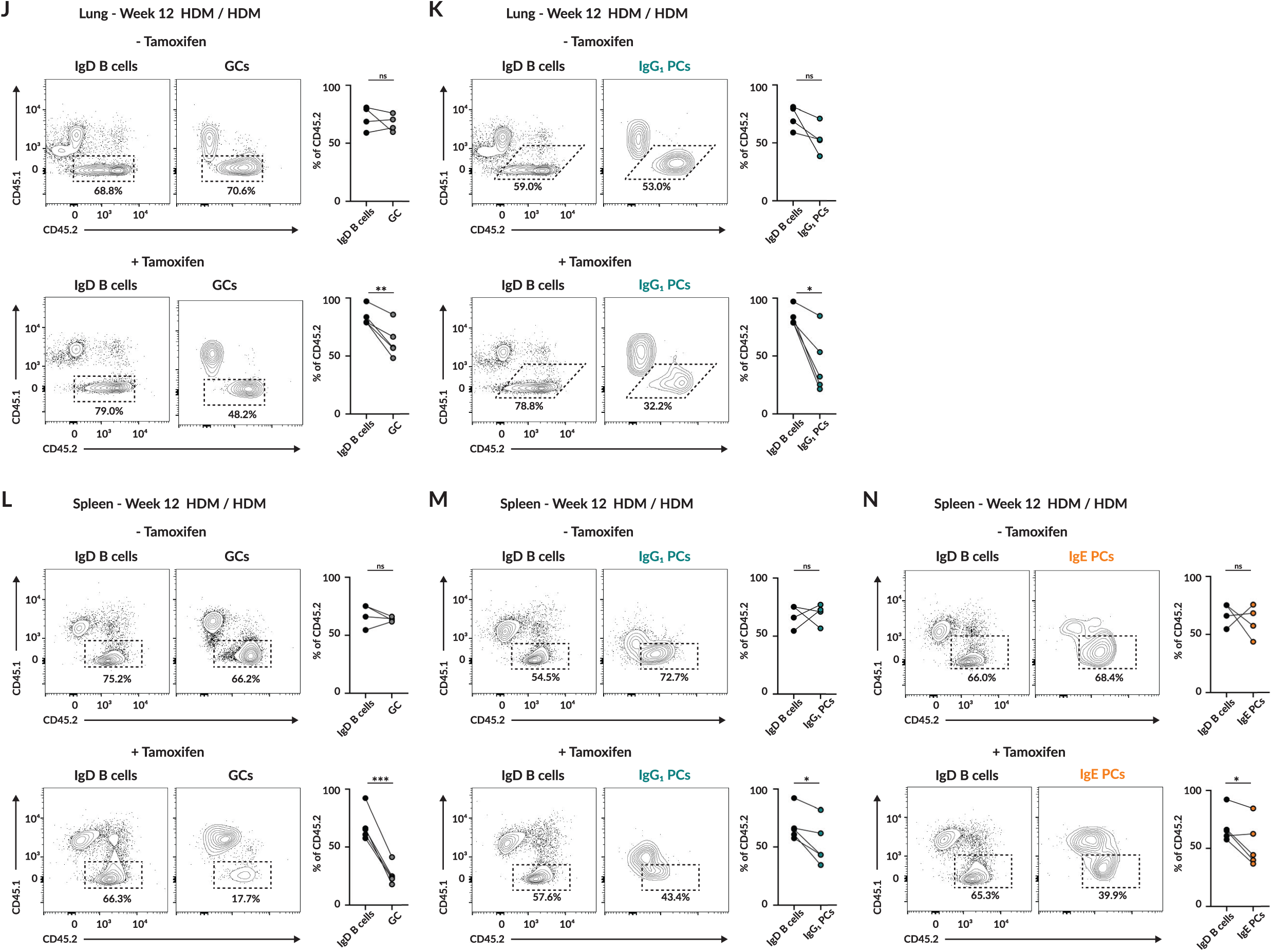
GC re-entry licenses MBCs for IgE production. (A) Contour plots display GC B cells in lung (top) and spleen (bottom) from mice treated with HDM extract and given anti-CD40L or left untreated as indicated in Figure 4G including enumeration of absolute cell number for each organ. (B) Contour plots show IgG_1_ and IgE PCs in lungs (top) and spleen (bottom) of mice following HDM treatment and given anti-CD40L or left untreated as depicted on Figure 4G with cell number quantification for each isotype and organ. (C) Schematic representation of experimental set up for HDM treatment and single dose anti-CD40L administration. (D) Contour plots display GC B cells in LN of mice immunized with HDM extract and given or not a single dose of anti-CD40L. (E) Contour plots showing IgG_1_ MBCs in LN of mice immunized with HDM extract and given or not a single dose of anti-CD40L. (F) Contour plots exhibit GC B cells in LN of mice immunized with HDM extract and given or not a single dose of anti-CD40L including quantification of the frequency of GC B cells. n=3. (G) Contour plots display IgG_1_ and IgE GC B cells in LN of mice immunized with HDM extract and given or not a single dose of anti-CD40L along with the enumeration of their frequency in the GC compartment. n=3. (H) Contour plots show GC B cells in lung (top) and spleen (bottom) of CD23Cre ^−/−^- and CD23Cre ^+/−^- Bcl6^flox^ mice treated with HDM extract as indicated on Figure 4K including enumeration of absolute cell numbers for each organ. (I) Contour plots exhibit IgG_1_ and IgE PCs in lung (top) and spleen (bottom) of CD23Cre ^−/−^-and CD23Cre ^+/−^- Bcl6^flox^ mice following HDM treatment outlined in Figure 4K including absolute cell number quantification for each isotype and organ. (J-K) Contour plots display CD45.1 and CD45.2 frequency in (J) IgD^+^ (left) and GC (right) and (K) within IgD^+^ (left) and IgG_1_ PC (right) compartments in lungs of chimeric mice reconstituted and treated as shown in figure 4O. (L-N) Contour plots show the CD45.1 and CD45.2 composition of (L) IgD^+^ (left) and GC (right), (M) IgD^+^ (left) and IgG_1_ PC (right) compartments and (N) IgD^+^ B cells (left) and IgE PCs (right) in spleens of chimeric mice built and subjected to treatments as indicated in Figure 4O. In all panels, quantification displays one representative experiment out of three.. Each dot corresponds to one mouse and bars represent mean ± s.e.m. We used non-parametric t-tests (A, B, H and I), one-way ANOVA (F and G), as well as paired t-tests (J-N) for statistical analysis. ns, non-significant; * p<0.05; ** p<0.01 and *** p<0.001.

**Figure S5.**
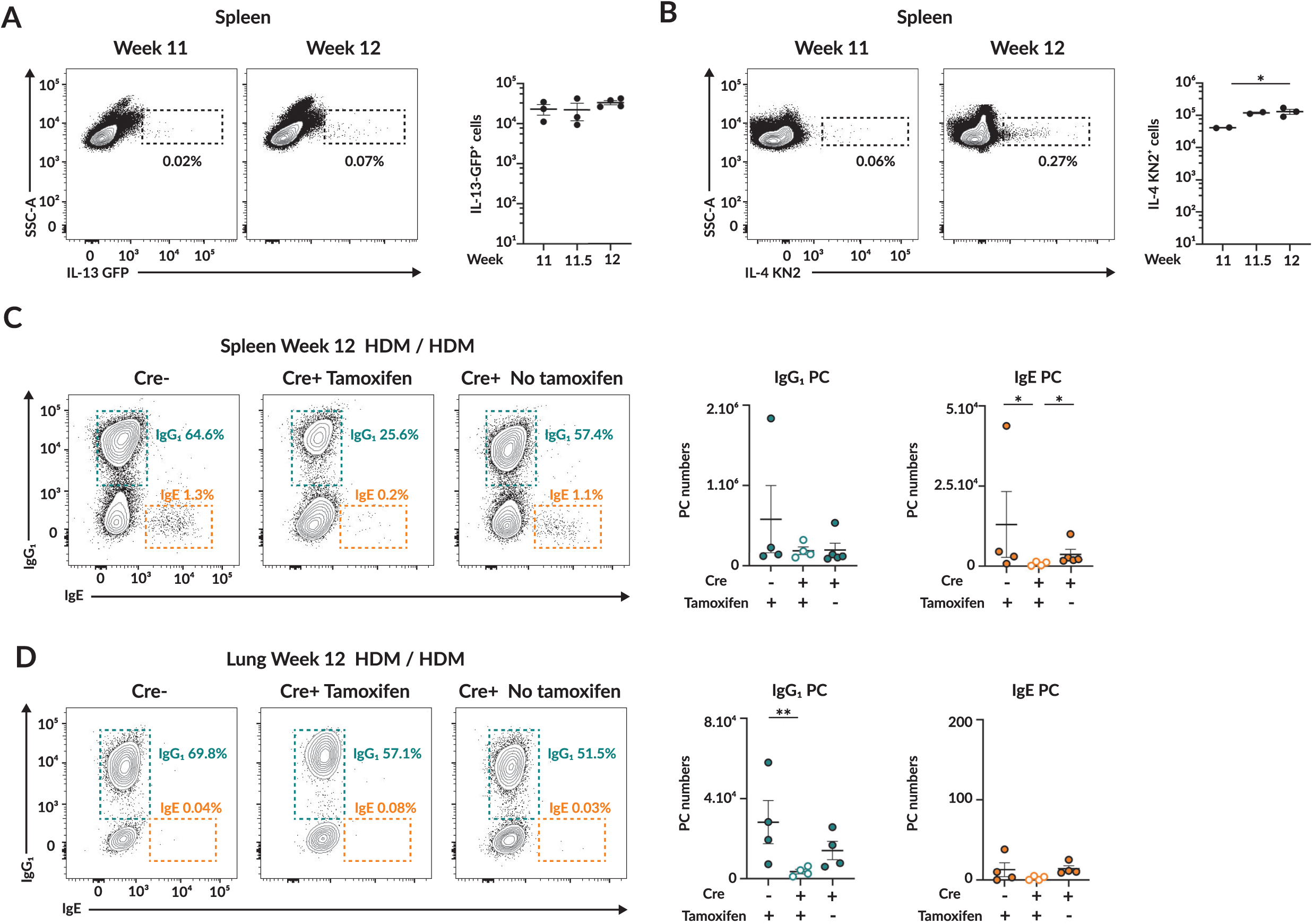
GC IL-4 microniches license MBCs for IgE differentiation. (A-B) Contour plots display (A) IL-13 GFP and (B) IL-4 KN2 staining in live spleen cells of IL-13 GFP and IL-4 KN2 mice respectively following treatment depicted in Figure 5D. (C-D) Contour plots show IgG1 and IgE PCs in (C) spleen and (D) lungs of Mb1Cre^ERT2^ IL4rα^flox^ mice treated as outlined in Figure 5I. In all panels, quantification displays one representative experiment out of three. Each dot corresponds to one mouse. We used non-parametric multiple t-tests for statistical analysis. * p<0.05 and ** p<0.01.

**Figure S6.**
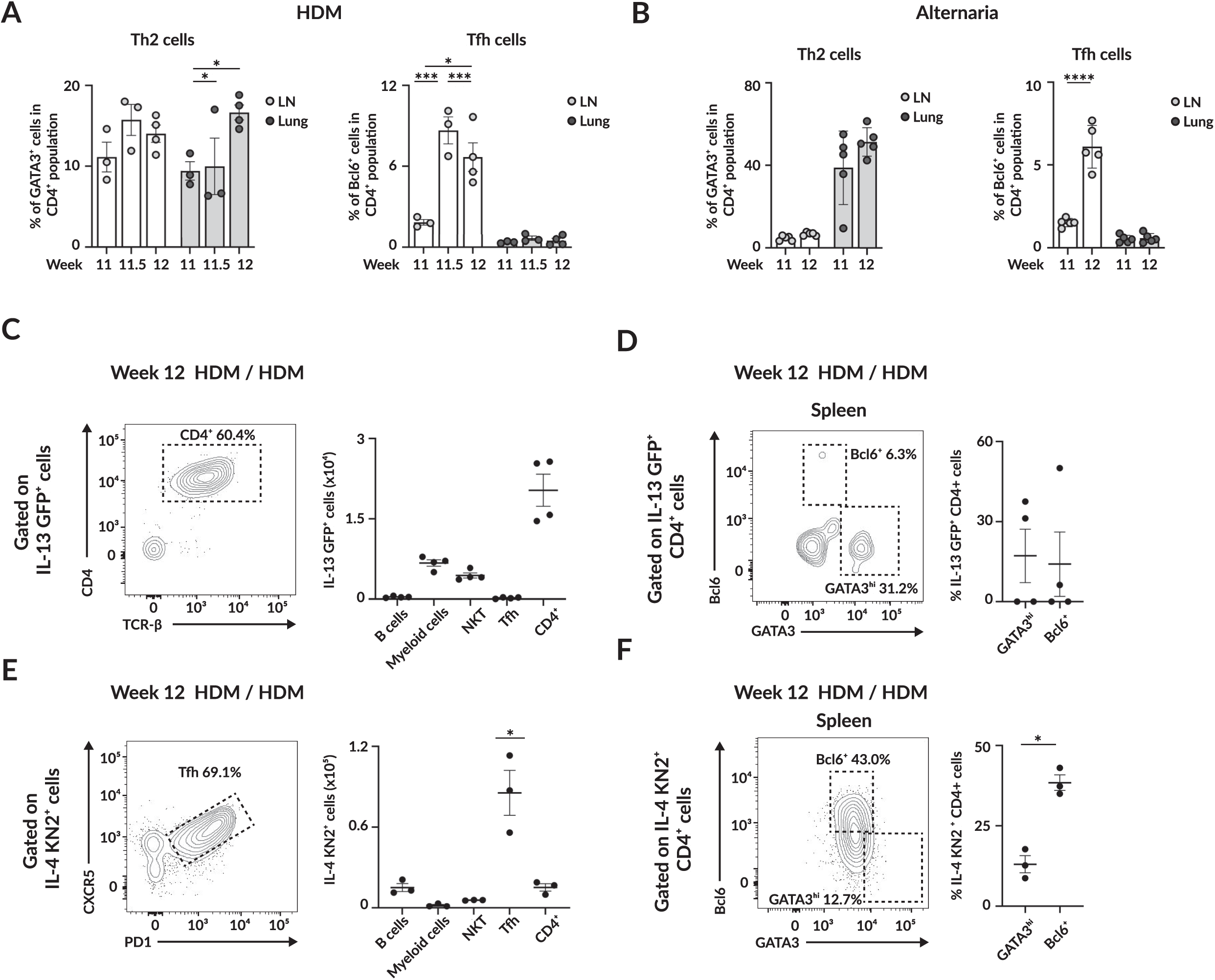
Tfh cells form GC IL-4 microniches essential for IgE recall. (A) Frequency of Th2 cells defined as GATA3^hi^ (left) and Tfh gated Bcl6^+^ (right) within CD4^+^ T cell compartment in LN and lungs of mice subjected to HDM treatment indicated in Figure 6A. (B) Th2 (gated as GATA3^hi^, left) and Tfh (defined as Bcl6^+^, right) frequency within CD4^+^ T cells in LN and lungs of mice following exposure to *Alternaria alternata* as shown in Figure 6D. (C) Contour plot show CD4 and TCR-o staining within IL-13 GFP^+^ cells in spleen of IL-13-GFP mice exposed to HDM extract as depicted in Figure 6F. (D) Contour plot display GATA3 and Bcl6 staining in CD4^+^ T cells and IL-13-GFP^+^ expressing cells in the spleen of mice subjected to HDM treatment indicated in FIgure 6F along with quantification of the frequency of expression of each transcription factor. (E) Contour plot exhibit CXCR5 and PD-1 staining in IL-4 KN2^+^ expressing cells in the spleen of IL-4 KN2 mice treated with HDM extract as shown in Figure 6F. (F) Contour plot displays GATA3 and Bcl6 staining within CD4^+^ and IL-4KN2^+^ expressing cells in the spleen of mice following exposure to HDM as indicated in Figure 6F along with the quantification of the expression of each transcription factor. In all panels, quantification displays one representative experiment out of three. Each dot corresponds to one mouse and bars represent mean ± s.e.m. We used non parametric multiple tests (A and B), one-way ANOVA (C and E) as well as paired t-tests (D and F) for statistical analysis. ns, non-significant; * p<0.05 and **** p<0.001.

## Material and methods

### Mice

Eight week-old C57BL/6 and BALB/c mice were purchased from Janvier Labs. IgEtdtTomato mice were generated by inserting a Tomato reporter gene linked to the second membrane exon of IgE by a T2A sequence in collaboration with Taconic Biosciences GmbH, Leverkusen, Germany. AidCre^ERT2^ mice were obtained from Claude-Agnès Reynaud and Jean Claude Weill, Institut Necker Enfants Malades, France. CD23^Cre^-Bcl6^flox^ mice were obtained from Michelle Linterman, Babraham Institute, Cambridge, United Kingdom. Rosa^eYFP^, CD45.1 and Mb1Cre^ERT2^ mice were purchased from Jackson Laboratories. μMT mice were obtained from Stéphane Mancini, Université de Rennes, France. Bcl6^flox^ mice were obtained from Nicolas Fazilleau, Infinity, Toulouse, France. We obtained IL-13 GFP mice and IL-13KO bone marrow cells from Judi Allen, The University of Manchester, Manchester, United Kingdom. IL-4 KN2 mice were obtained from Andrew McDonald, The University of Edinburgh, Edinburgh, United Kingdom. RAG2KO and IL4rα^flox^ mice were obtained from Toby Lawrence, Centre d’Immunologie Marseille-Luminy, Marseille, France. IL-4KO bone marrow cells were obtained from Adriana Gruppi, Centro de Investigación Bioquímica Clínica e Inmunología, Córdoba, Argentina. Bone marrow cells from CD4Cre^ERT2^-Bcl6^flox^ mice were obtained from Carolyn King, University of Basel, Basel, Switzerland. AidCre^ERT2^ mice were further crossed with Rosa^eYFP^, whereas Mb1Cre^ERT2^ mice were also crossed with Bcl6^flox^ and IL4rα^flox^ respectively.

To generate mixed bone marrow chimeras, μMT mice of 6-8 weeks of age were irradiated with 2 doses of 4.75 Gy, 4 hours apart, whereas 6-8 week-old RAG2KO mice were irradiated with 2 doses of 3 Gy, 4 hours apart. The day after, we i.v. injected bone marrow cells (5×10^6^ cells in total) in recipient mice. Injected animals were kept under Bactrim, administered in water, for 3 days prior and 3 weeks post irradiation. Chimeras were used after 8 weeks of reconstitution.

Mice were bred and kept in the animal facilities at the Centre d’Immunologie Marseille-Luminy (CIML). A maximum of 5 mice were housed in each cage under a 12 hour light/dark cycle at a constant temperature of 22°C (19°C-23°C temperature range). Mice were fed with autoclaved standard chow diet and reverse osmosis water. Each cage contained 5 mm of aspen chip and tissue wipes for bedding and a cellulose mouse dome to enrich mice’ environment. Eight to twelve week-old mice were used for experimentation and littermates (males and females) were randomly assigned to experiments in groups of 3 to 6 mice. All experiments were carried out according to French and European guidelines for laboratory animal welfare and under the animal license 2023072010331744 reviewed and approved by the local animal ethics committee in Marseille.

### Immunisation and injections

Mice were subjected to light isoflurane anaesthesia (1.5 to 3% isoflurane) prior to each intranasal administration of 40 μg of *Dermatophagoides pteronyssinus* extract (HDM, CiteQ Biologics) freshly reconstituted in 40 μl of sterile PBS 1X. Mice exposed to Alternaria alternata were given intranasally 40 μg of *Alternaria alternata* medium (*Alt.*, CiteQ Biologics) freshly reconstituted in 40 μl of sterile PBS 1X. AidCre^ERT2^ -Rosa^eYFP^, Mb1Cre^ERT2^-IL4rα^flox^ mice as well as CD45.1/Mb1Cre^ERT2^-Bcl6^flox^, CD45.1/Mb1Cre^ERT2^-IL4rα^flox^ and CD4Cre^ERT2^-chimeric mice received three doses of 5 mg tamoxifen (Cayman chemical) resuspended in 100 μl sterile corn oil (Sigma Aldrich) by gavage followed by maintenance in tamoxifen diet (0.5 g per kg tamoxifen with 5% saccharose, 25kGY irradiated, Safe Custom Diets) for 5 weeks for AidCre^ERT2^ -Rosa^eYFP^ and until the end of the experiment for all other mice and chimeras. To block the egress of immune cells, mice were given 2.5 μg/mL FTY720 (Sigma Aldrich) in their drinking water, which was kept in opaque bottles and replaced every two days. To disrupt GCs *in vivo*, mice were injected with either 300 μg of anti-CD40L (clone MR1, BioXCell) or respective isotype control (*InVivo*Mab polyclonal Armenian Hamster IgG, BioXCell) in 200 μl of sterile PBS at different time points during HDM rechallenge. To block IL-4 or IL-13 *in vivo* upon HDM re-exposure, 150 μg of anti-IL-4 (clone 11B11, BioXcell) or 150 μg of anti-IL-13 (clone eBio1316H, ThermoFisher Scientific) in 200 μl of sterile PBS were administered to mice at different days of rechallenge. To label immune cells in circulation, 5 μg of anti-CD45 antibody was injected i.v. 5 minutes prior to sacrifice. Bone marrow cells (5×10^6^) were resuspended in 100 μl of sterile PBS and injected i.v.

| Reagent/Antibodies | Source | Catalog |
| --- | --- | --- |
| *Dermatophagoides pteronyssinus* extract | CiteQ Biologics | 02.01.85, Lot 20B12 |
| *Alternaria alternata* medium | CiteQ Biologics | 09.01.26, Lot 18L17 |
| Tamoxifen | Cayman Chemical | 10540-29-1 |
| Corn oil | Sigma Aldrich | C8267 |
| Tamoxifen pellets 0.5g/kg tamoxifen + 5% saccharose 25kGy | Safe diets | E8220 |
| FTY720 | Sigma Aldrich | SML0700 |
| Purified Armenian Hamster Anti-Mouse CD40L, clone MR1 | BioXcell | BE0017 |
| *InVivoMab* polyclonal Armenian Hamster IgG | BioXcell | BE0091 |
| Purified Anti-Mouse IL-13, clone eBio1316H | ThermoFisher Scientific | 16-7135-85 |
| Purified Anti-Mouse IL-4, clone 11B11 | BioXcell | BE0045 |
| Biotinylated Rat Anti-Mouse CD45, clone 30-F11 | Biolegend | 103104 |

### Flow cytometry

Single cell suspension of mLN and spleen were obtained by smashing organs through a 70 μm nylon filter with the aid of a plastic plunge in 2% FCS in PBS. To obtain a lung single cell suspension, lungs were digested using the lung dissociation kit (Milteny) according manufacturer instructions. Lung single cell suspension was further enriched for its immune cell fraction using a 40 to 80% Percoll (GE Healthcare) gradient. Spleen and lung cell suspensions were then incubated with 1 mL of red blood cell lysis buffer (invitrogen) for 5 minutes at room temperature. To block non-specific binding and label dead cells, organ cell suspensions were incubated with hybridoma supernatant 2.4G2 and Live/Dead fixable blue viability kit (ThermoFisher Scientific) used according to manufacturer instructions in 2% FCS in PBS for 20 minutes on ice. Surface cell markers were labelled with the indicated anti-mouse antibodies for 20 minutes on ice. Then, cell suspensions were fixed, permeabilised and stained targets intracellularly using the BD Cytofix/Cytoperm Fixation and Permeabilisation solution (BD Biosciences) according to manufacturer instructions. Unlabelled anti-mouse IgE (clone R35-72, BD Biosciences) was used to saturate surface bound IgE followed by intracellular staining with the same anti-IgE clone to label IgE bearing B cells as previously described^42^. Alternatively, to stain for transcription factors, single cell suspensions were fixed, permeabilised and stained with antibodies targeting mouse transcription factors using Foxp3/Transcription factor staining buffer set (ThermoFisher Scientific) according to manufacturer instructions. Finally, cells were resuspended in 400 μl 2% FCS in PBS along with counting beads CountBright (ThermoFisher Scientific) and analyzed on Fortessa X20/Symphony A5/LSR II UV (BD Bioscience). When sorting cells, single cell suspensions were not fixed and DAPI (0.1 μg/mL final concentration) was added right before sample acquisition to exclude dead cells. Sorting was carried out in FACSAria II (BD Biosciences). Analysis of flow cytometry data was carried out using Flowjo (BD Biosciences) software version 10.

| **Antibody** | **Source** | **Catalog** |
| --- | --- | --- |
| Purified Rat Anti-Mouse IgE, clone R35-72 | BD Biosciences | 553413 |
| Alexa Fluor 488 Mouse Anti-Mouse/Human GATA3, clone L50-823 | BD Biosciences | 560163 |
| Alexa Fluor 488 Rat Anti-Mouse IgD, clone 11-26c.2a | Biolegend | 405718 |
| Alexa Fluor 647 Mouse Anti-Mouse/Human Bcl6, clone K112-91 | Biolegend | 561525 |
| Alexa Fluor 647 Rat Anti-Mouse/Human GL7, clone GL7 | Biolegend | 144606 |
| Alexa Fluor 700 Rat Anti-Mouse CD38, clone 90 | Biolegend | 560381 |
| APC CD1d tetramer | NIH tetramer core facility | N/A |
| APC Fire 750 Rat Anti-Mouse PD-1 (CD279), clone 29T.1A12 | Biolegend | 1352540 |
| APC/Cy7 Rat Anti-Mouse CD19, clone 6D5 | Biolegend | 115530 |
| BUV395 Rat Anti-Mouse CD38, clone 90 | BD Biosciences | 740245 |
| BUV395 Mouse Anti-Mouse/Human GATA3, clone L50-823 | BD Biosciences | 565448 |
| BUV737 Rat Anti-Mouse CD93, clone AA4.1 | BD Biosciences | 741800 |
| BUV737 Rat Anti-Mouse B220 (CD45R), clone RA3-6B2 | BD Biosciences | 563793 |
| BUV737 Mouse Anti-Mouse CD45.2, clone 104 | BD Biosciences | 612778 |
| BV421 Rat Anti-Mouse CXCR5 (CD185), clone L138D7 | Biolegend | 145511 |
| BV510 Rat Anti-Mouse IgE, clone R35-72 | BD Biosciences | 747734 |
| BV510 Armenian Hamster Anti-Mouse Fas, clone Jo2 | BD Biosciences | 563646 |
| BV605 Rat Anti-Mouse CD4, clone GK1.5 | Biolegend | 100451 |
| BV605 Rat Anti-Mouse CD45, clone 30-F11 | Biolegend | 4060451 |
| BV650 Rat Anti-Mouse CD93, clone AA4.1 | BD Biosciences | 752943 |
| BV650 Armenian Hamster Anti-Mouse TCR-α, clone H57-597 | Biolegend | 109251 |
| BV711 Rat Anti-Mouse CD138, clone 281-2 | Biolegend | 142519 |
| BV785 Rat Anti-Mouse IgD, clone 11-26c.2a | BD Biosciences | 563618 |
| BV785 Rat Anti-Mouse CD138, clone 281-2 | Biolegend | 142534 |
| BV785 Mouse Anti-Human CD2, clone RPA-2.10 | Biolegend | 300233 |
| FITC Rat Anti-Mouse IgE, clone R35-72 | BD Biosciences | 553415 |
| FITC Rat Anti-Mouse CD45, clone 30-F11 | Biolegend | 103107 |
| FITC Mouse Anti-Human CD2, clone RPA-2.10 | Biolegend | 300206 |
| Pacific Blue Rat Anti-Mouse/Human GL7, clone GL7 | Biolegend | 144613 |
| PE Rat Anti-Mouse CD45, clone 30-F11 | Biolegend | 103105 |
| PE Rat Anti-Mouse CD98, clone RL388 | Biolegend | 128208 |
| PE Rat Anti-Mouse CXCR5 (CD185), clone L138D7 | Biolegend | 145503 |
| PE/Cy5 Rat Anti-Mouse CD19, clone 6D5 | Biolegend | 115510 |
| PE/Cy7 Rat Anti-Mouse CD39, clone 24DMS1 | ThermoFisher Scientific | 250391 |
| PE/Cy7 Rat Anti-Mouse CD4, clone GK1.5 | Biolegend | 100421 |
| PE/Cy7Rat Anti-Mouse IgM, clone R6-60.2 | BD Biosciences | 552867 |
| PerCP/Cyanine5.5 Rat Anti-Mouse IgM, Clone RMM-1 | Biolegend | 406512 |
| PerCP/Cyanine5.5 Rat Anti-Mouse/Human GL7, clone GL7 | Biolegend | 144610 |
| PerCP/Cyanine5.5 Mouse Anti-Mouse CD45.1, clone A20 | Biolegend | 110728 |
| V450 Rat Anti-Mouse IgG1, clone A85-1 | BD Biosciences | 562107 |
| Biotinylated Rat Anti-Mouse CD45, clone 30-F11 | Biolegend | 103104 |
| AF488 Streptavidin | Biolegend | 405235 |
| BV650 Streptavidin | Biolegend | 405232 |
| PE Streptavidin | ThermoFisher Scientific | S866 |
| APC/Cy7 Streptavidin | Biolegend | 405208 |

### Immunohistochemistry

LNs and lungs were fixed in 4% PFA (Electron Microscopy Sciences) for 3 hours at 4°C followed by a quick wash in PBS and further incubation in 30% sucrose in PBS overnight at 4°C. Then, organs were embedded in OCT solution and snapped-frozen in liquid nitrogen-cooled isopentane. Alternatively, lymph nodes and lungs of IL-4 KN2 mice were freshly frozen in OCT solution as indicated above without prior fixation. Cryostat sections (20 μm) were dried in silica beads or fixed with ice-cold acetone for 10 minutes for organ sections of IL-4 KN2 mice followed by 2 washes in PBS. Then, sections were blocked in 2% BSA, 1% FCS, 1% goat serum and 0.5% saponin in PBS for 45 minutes at room temperature. Following blocking, sections were incubated with primary antibodies in 2% BSA, 1% FCS, 1% goat serum and 0.5% saponin in PBS for 3 hours at room temperature, washed and further incubated with secondary antibodies for 3 more hours at room temperature or overnight at 4°C. To label Bcl6, sections stained with primary antibodies were fixed and permeabilised using Foxp3/Transcription factor staining buffer set (ThermoFisher Scientific) prepared according to manufacturer instructions followed by blocking in 2% BSA, 1% FCS and 1% goat serum in FoxP3 staining kit permeabilisation solution 1x for 1 hour at room temperature. Then, sections were incubated with secondary antibodies including Bcl6 in 2% BSA, 1% FCS and 1% goat serum in FoxP3 staining kit permeabilisation solution 1x overnight at 4°C. Lastly, sections were washed in PBS and mounted in Fluoromont-G mounting media (Invitrogen). Images were acquired in a LSM 880 AiryScan (Zeiss) inverted confocal microscope using a Plan-Apochromat 10x/0.45 for whole organ imaging or Plan-Apochromat 20x/0.8.

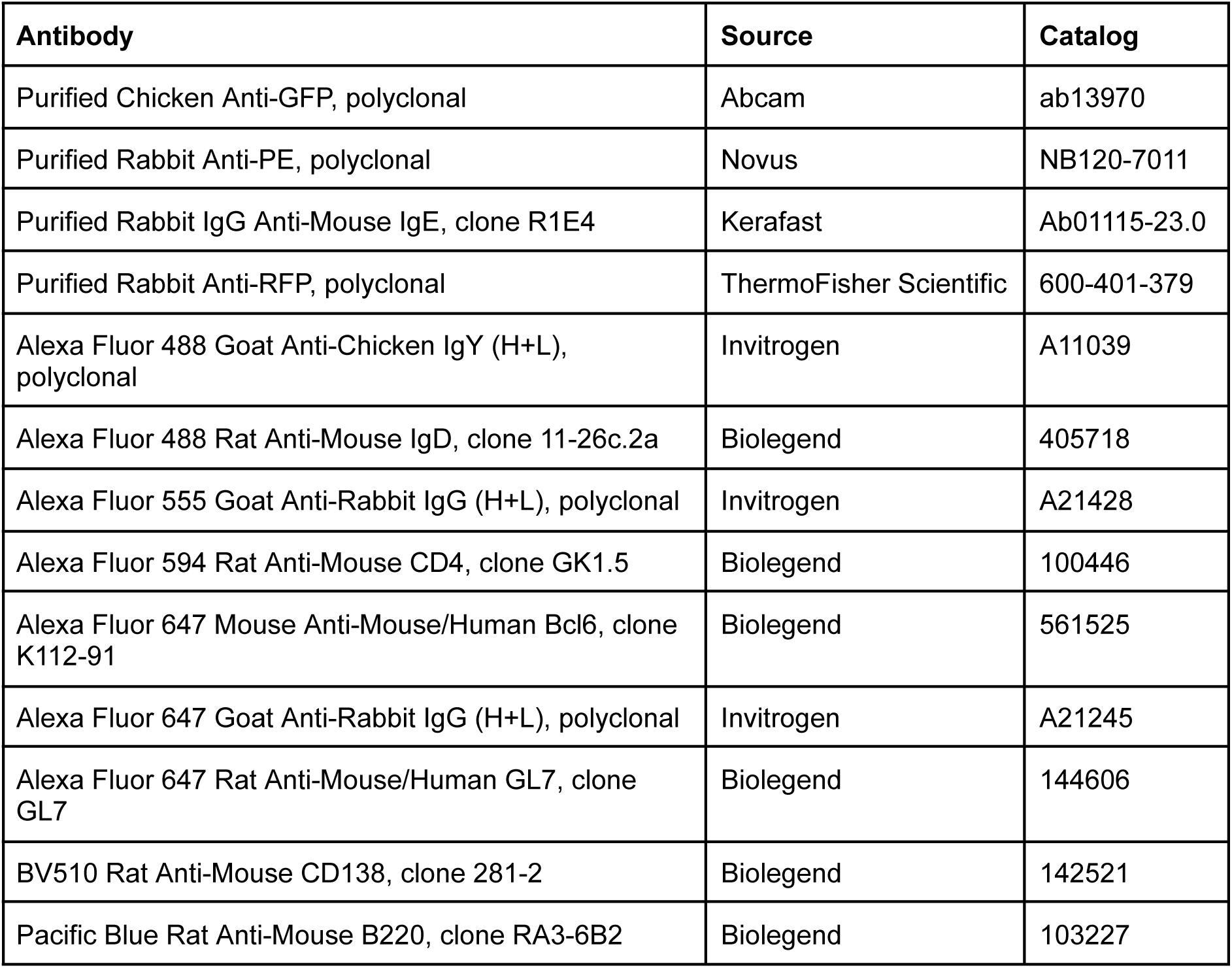
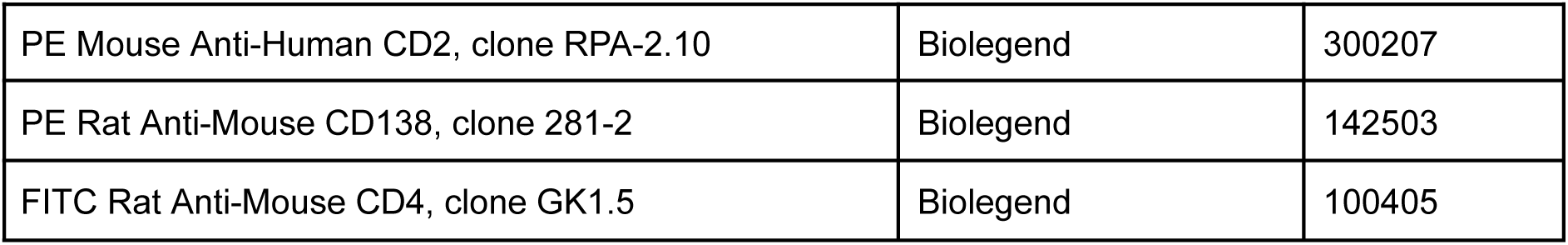

### *Ex vivo* culture of MBCs

Sorted LN and lung MBCs were cultured in U-bottom 96 well plates in the presence of 1 μg/mL recombinant mouse CD40l/TNFSF5 (HA-Tag) (R&D Systems), 50 ng/mL recombinant mouse IL-4 (Prepotech) and 10 ng/mLrecombinant IL-5 (Prepotech) in 10% FCS, 100 u/mL penicillin, 100 μg/mL streptomycin, 1mM sodium pyruvate and 5×10^−5^ M beta-mercaptoethanol in RPMI medium for 96 hours at 37°C and 5% CO_2_. At indicated time points, cells were collected, stained and analyzed by flow cytometry.

| Chemical | Source | Catalog |
| --- | --- | --- |
| Recombinant mouse CD40l/TNFSF5 (HA-Tag) | R&D | 8230 |
| Recombinant mouse IL-4 | Prepotech | 214-14 |
| Recombinant mouse IL-5 | Prepotech | 215-15 |

### ELISA

To measure total IgE and Der p 1 specific antibodies, ELISA (enzyme-linked immunoabsorbent assay) plates were either coated with 2 μg/mL unlabelled anti-mouse IgE (clone R35-72, BD Biosciences) to measure total IgE and Der p 1 IgE or with 2 μg/mL recombinant Der p 1 (CiteQ Biologics) to assess Der p 1 IgG_1_ titers in carbonate buffer (pH=9.6) overnight at 4°C. Then, plates were washed with 0.01%Tween in PBS (PBS-T) followed by blocking in 2% BSA in PBS-T for 3 hours at room temperature and incubation with serial dilutions of serum supernatants in 0.5% BSA in PBS-T overnight at 4°C. The following day, plates were washed with PBS-T and probed for 2 hours at room temperature with 1 μg/mL anti-mouse IgE-biotin (Southern Biotech) for total IgE detection, Der p 1-biotin (CiteQ Biologics) for Der p 1 IgE estimation or anti-mouse IgG_1_-biotin (Southern Biotech) for the measurement of Der p 1 IgG_1_. Then, plates were washed with PBS-T and incubated with 0.2 μg/mL Streptavidin-Alkaline Phosphatase (Sigma Aldrich). Finally, plates were washed with PBS-T and then incubated with freshly prepared p-NitrophenylPhosphate (Sigma Aldrich) solution. Absorbance was measured at 405 nm using a SPECSTROstar Omega (BMG Labtech) plate reader.

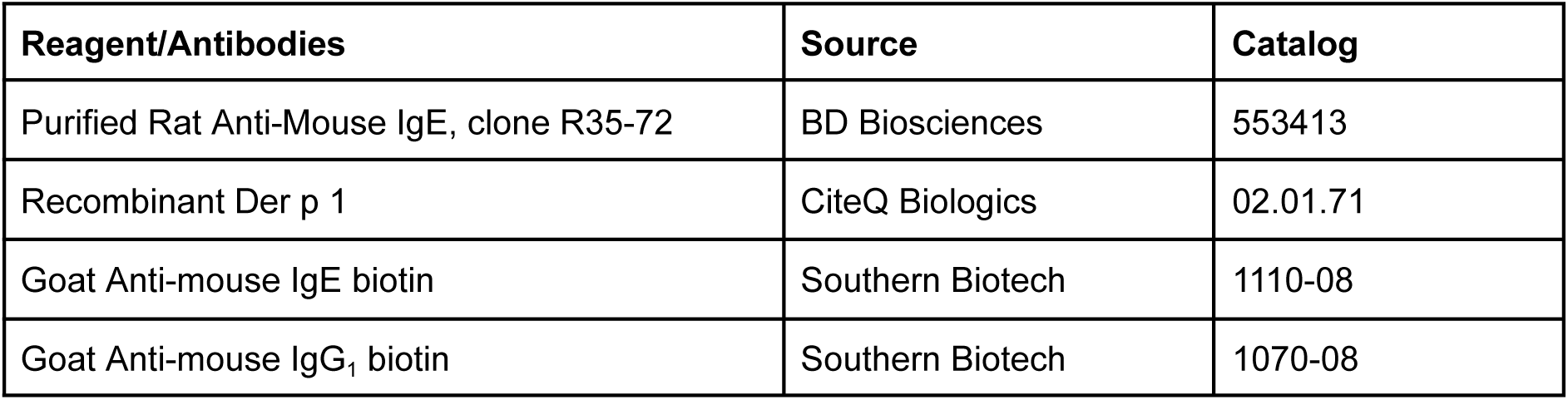
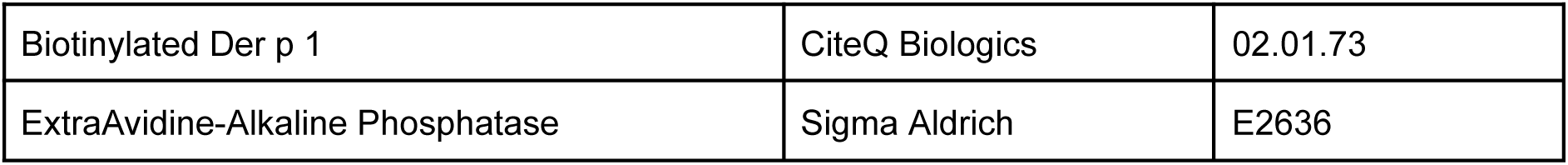

| Reagent/Antibodies | Source | Catalog |
| --- | --- | --- |
| Purified Rat Anti-Mouse IgE, clone R35-72 | BD Biosciences | 553413 |
| Recombinant Der p 1 | CiteQ Biologics | 02.01.71 |
| Goat Anti-mouse IgE biotin | Southern Biotech | 1110-08 |
| Goat Anti-mouse IgG<sub>1</sub> biotin | Southern Biotech | 1070-08 |
| Biotinylated Der p 1 | CiteQ Biologics | 02.01.73 |
| ExtraAvidine-Alkaline Phosphatase | Sigma Aldrich | E2636 |

### 10x 5’ scRNA-seq library preparation

Both mLNs and lungs of AidCre^ERT2^-Rosa^eYFP^were digested to obtain a single cell suspension using lung dissociation kit (Milteny) according manufacturer instructions to avoid batch effects resulting from the use of different approaches to obtain organ cell suspensions^61^. Single cell suspensions were filtered twice through a 70 μm nylon mesh and then blocked and stained with cell surface antibodies as indicated above in section Flow cytometry. Then, single cell suspensions from each organ were independently barcoded with hashtags containing a mixture of two specific antibodies for CD45 and MHC I (Biolegend) in 2% FCS in PBS for 30 minutes on ice followed by a wash and resuspension in 500 μl of 2% FCS in PBS. DAPI (0.1 μg/mL final concentration) was added right before sample acquisition to exclude dead cells. For each sample up to 5000 live YFP+ IgD^−^ CD45 i.v.^−^ PCs (CD39^+^ CD98^+^) and up to 8000 live YFP+ IgD^−^ CD45 i.v.^−^ CD39^−^ CD98^−^ B cells for each mouse and organ were bulk sorted with BD FACS Aria II. Sorted PCs as well as sorted non-PCs were pooled respectively and loaded in two single capture wells for downstream 10x Genomics Single cell 5’ workflow. To account for differential gene expression between PCs and non-PCs, separate libraries were built for these two populations.

10x 5’ scRNA-seq libraries were prepared following instructions from the manufacturer with minor modifications for the generation of BCR-seq libraries. Right after cDNA amplification, the large cDNA fraction originated from cellular mRNAs was separated from the Hashtag (HTO)-containing fraction using SPRI beads. Fifty and ten ng of mRNA-derived cDNA were used to generate the transcriptome and the BCR library respectively. Gene expression libraries were prepared following manufacturer instructions, while the BCR library was built by amplifying heavy and light chains by two rounds of PCR (6 cycles + 8 cycles) using external primers recommended by 10x Genomics. Eight hundred pg of amplified BCR cDNA were fragmented using a Nextera XT DNA primer (10x Genomics) and a Nextera i7 reverse primer (Illumina). Five ng of the HTO containing cDNA fraction were used to produce the HTO library. All three resulting libraries from PCs and non-PCs were pooled together and sequenced in an Illumina NextSeq550 platform using High Output 75 cycle flow cells, targeting 5×10^4^ reads/cell for gene expression, 5×10^3^ reads/cell for BCR and 2×10^3^ reads/cell for HTO in paired-end single-index mode.

### scRNA and BCR-seq analysis

Read count-UMI matrices were generated from FASTQ files with Cell Ranger (7.0.1), and then imported into R using the Seurat package^62^. Out of 12121 cells, a cluster of 494 cells identified as T-cell contamination was marked for removal in downstream analyses. Seurat function HTODemux was used to demultiplex individual mice, and tissue, after ‘CLR’ normalization of HTO counts. Gene expression values were normalized by total UMI counts per cell, multiplied by 10000 and log10 transformed. Highly variable genes were identified using the “FindVariableFeatures” function in Seurat and linear transformed prior to dimensionality reduction by principal component analysis (PCA). To enable cell types identification independently of differential immunoglobulin expression by dimensionality reduction and unsupervised clustering approaches, genes coding for heavy and light chain immunoglobulins were individually excluded from the list of variable genes. Clustering and non-linear dimensionality reduction was performed using Uniform Manifold Approximation and Projection (UMAP) with 50 principal components and variable resolution to identify cell populations based on gene expression. Marker genes were identified with the Seurat function “FindAllMarkers” (wilcoxon). A transcriptome dynamic analysis was performed using scVelo^63^ after computing spliced, unspliced, and ambiguous count matrices with Velocito^64^. These matrices were injected into the Seurat object, which has been converted to Anndata for scVelo analysis in python. Genes list imported from Cui et al, 2024 were grouped by cytokine/molecule, the Seurat function AddModuleScore was used to compute an enrichment score based on corresponding list of genes^33^. Resulting scores were represented as violin-plots (grouped by previously identified clusters), and as quantitative features on UMAP.

To perform BCR analyses, the filtered cell Ranger output was imported by package scRepertoire^65,66^ to combine heavy and light chain sequences into clones using the ‘strict’ definition of a clone: heavy chain VDJ gene segment usage and complementarity determining region 3 (CDR3) nucleotide sequence. BCR information was then combined to gene expression in the Seurat object for each mouse individually. Clonal overlap analyses between tissues and cell subsets were visualized using the circlize package^67^. The clonalNetwork function from scRepertoire was used to generate a network based on clonal proportions of cell identities, and superimpose the network onto precomputed UMAP.

### Statistical analysis

Statistical parameters including the exact value of n (number of individuals), the mean, the standard error of the mean (s.e.m.), the p values and the statistical test used are stated in each figure and figure legends. Statistical analyses were carried out using GraphPad Prism 10 (GraphPad Software) including t-tests, multiple t-tests, paired t-tests, one-way and two-way ANOVA. Turkey post-hoc tests were used for correcting for multiple comparisons in ANOVA tests. Any p value below 0.05 was considered significant. In figures, ns stands for non-significant and asterisks represent: * p<0.05; ** p<0.01; *** p<0.001 and **** p<0.001.

